# Developmental mechanisms contributing to non-linear firing dynamics in spinal motoneurons of the postnatal mouse

**DOI:** 10.64898/2026.03.12.711366

**Authors:** Simon A. Sharples, Gareth B. Miles

## Abstract

The intrinsic properties of spinal motoneurons support flexible movement, including the maintenance of postural tone. Motoneurons can produce sustained action potential output that outlasts synaptic input, a phenomenon traditionally attributed to persistent inward currents (PICs) mediated by sodium and calcium channels. Using whole-cell patch clamp electrophysiology, we examined how specific ion channels contribute to PIC maturation and non-linear firing dynamics that allow motoneurons to sustain their output in fast and slow lumbar motoneurons across postnatal development in mice. PIC amplitude and non-linear firing dynamics increased after weight bearing in fast but not slow motoneurons. Blocking Nav1.6 channels reduced PIC amplitude at both pre- and post-weight-bearing stages, whereas L-type calcium channel blockade only reduced PICs after weight bearing emerged. However, reducing PIC amplitude—either individually or in combination—did not abolish sustained firing hysteresis. Unexpectedly, activation of muscarinic receptors increased PIC amplitude while promoting adaptive firing dynamics, suggesting that PICs alone do not drive this behavior. Instead, pharmacological manipulation of potassium currents mediated by KCNQ and Kv1.2 channels, which oppose PICs, produced substantial changes in firing dynamics. Strikingly, blocking HCN channels promoted sustained firing dynamics and led to the emergence of self-sustained firing in fast motoneurons. These results indicate that while PICs and non-linear firing dynamics mature together, sustained firing relies on mechanisms beyond PICs, with potassium and HCN channels playing key modulatory roles.

**Key Points:**

- Persistent inward currents (PICs) and recruitment-derecruitment hysteresis increase in parallel in fast, but not slow, motoneurons following the onset of hindlimb weight bearing.
- Increased expression or function of L-type calcium channels may contribute to enhanced PICs in fast motoneurons after weight bearing emerges.
- Neither Nav1.6 nor L-type calcium channels are required for sustained firing hysteresis in fast motoneurons.
- KCNQ channels attenuate PICs and, together with Kv1.2 channels, shape recruitment–derecruitment asymmetry, thereby modulating firing hysteresis in fast motoneurons.
- HCN channels generate a resting H-current that delays recruitment, modulates firing hysteresis, and prevents the emergence of self-sustained firing in fast motoneurons.

## Introduction

The development of complex motor control is a fundamental aspect of early life, enabling animals to interact with their environment, maintain posture, and execute coordinated movements. In rodents, the emergence of hindlimb weight bearing represents a critical milestone in motor development, typically occurring during the second half of the second postnatal week (Altman & Sudarshan, 1975). This behavioral transition reflects the maturation of spinal circuits and the intrinsic properties of motoneurons (Vinay *et al*., 2000; Stifani, 2011; Quinlan *et al*., 2011; Durand *et al*., 2015; Jean-Xavier *et al*., 2018; Smith & Brownstone, 2020; Sharples & Miles, 2021), which must generate sustained activation of muscle fibers to support postural stability and locomotion (Heckman *et al*., 2008b).

Persistent inward currents (PICs) allow motoneurons to maintain firing once activated, which is believed to contribute to supporting postural tone. These currents, conducted by sodium (NaV1.6) and calcium (CaV1.3) channels, amplify excitatory inputs and help maintain the membrane potential above the threshold for action potential generation, thereby supporting sustained firing beyond the initial synaptic input (Schwindt & Crill, 1977, 1980; Hounsgaard & Kiehn, 1989; Carlin *et al*., 2000; Powers & Binder, 2003; Li & Bennett, 2003; Li *et al*., 2004; Heckmann *et al*., 2005; Elbasiouny *et al*., 2006; Harvey *et al*., 2006a; Manuel *et al*., 2009; Bouhadfane *et al*., 2013; Powers & Heckman, 2015; Mesquita *et al*., 2024). In other words, a brief excitatory signal can produce ongoing neuronal activity because the inward current “holds” the membrane potential near or above firing threshold. PICs also contribute to nonlinear membrane responses, meaning that the relationship between input and output is not strictly proportional—small changes in input can produce large changes in firing—and can create bistable firing patterns, in which a neuron can stably remain in either a low- or high-firing state depending on prior activity (Bos *et al*., 2018, 2021; Drouillas *et al*., 2023; Harris-Warrick *et al*., 2024).

A frequently studied feature associated with PICs is recruitment–derecruitment asymmetry (Bennett *et al*., 2001; Li & Bennett, 2003; Harvey *et al*., 2006b; Button, 2008; Manuel *et al*., 2009; Meehan *et al*., 2010b, 2010a; Quinlan *et al*., 2011; MacDonell *et al*., 2015; Huh *et al*., 2017). This is typically measured using a triangular current injection, where the current is gradually increased and then decreased. Motoneurons often begin firing at a higher current on the ascending limb of the ramp (recruitment) than the current at which firing stops on the descending limb (derecruitment), producing a characteristic “hysteresis” in the firing–current relationship (Bennett *et al*., 2001). In addition, PICs can support higher firing rates on the descending limb compared with the ascending limb, believed to be reflecting the sustained depolarizing influence of inward currents. Similar patterns are observed in human motor units during triangular isometric contractions (Gorassini *et al*., 2002; Udina *et al*., 2010), highlighting the translational relevance of these firing behaviors. While PICs are believed to be central to these phenomena, other intrinsic mechanisms—including spike-frequency adaptation, spike-threshold accommodation, and potassium conductances—can also influence recruitment, derecruitment, and firing rates independently of inward currents (Sawczuk *et al*., 1995; Button *et al*., 2007; Li & Bennett, 2007; Powers *et al*., 2008; Kalmar *et al*., 2009; Hamm *et al*., 2010; Revill & Fuglevand, 2011; Vandenberk & Kalmar, 2014; Powers & Heckman, 2015; Leroy *et al*., 2015; Bos *et al*., 2018; Sharples *et al*., 2023; Deutsch & Elbasiouny, 2024; Molkov *et al*., 2025). Together, these mechanisms suggest that sustained firing and hysteresis arise from a dynamic balance of inward and outward currents.

Previous work has emphasized the role of dendritic CaV1.3 currents in mediating hysteresis under strong neuromodulatory drive (Heckman *et al*., 2003, 2008a), but other ion channels at the axon initial segment (AIS) are also likely to play a critical role. NaV1.6 channels generate a persistent sodium current, while potassium channels such as Kv1.2 and KCNQ are positioned to shape action potential initiation and repetitive firing (Bos *et al*., 2018; Drouillas *et al*., 2023; Sharples *et al*., 2023; Harris-Warrick *et al*., 2024). These AIS potassium channels may not simply oppose PICs (Verneuil *et al*., 2020), but likely actively determine the timing of recruitment and derecruitment, offering a complementary mechanism that can influence firing hysteresis. Understanding the relative contributions of inward versus outward conductances to the maturation of motoneuron firing properties is especially important during the postnatal period when hindlimb weight bearing emerges, as disruptions in these mechanisms may underlie motor disorders or delayed motor development.

In this study, we investigated the maturation of PICs and recruitment–derecruitment asymmetry in fast and slow motoneurons across the postnatal period surrounding the onset of hindlimb weight bearing. Using whole-cell patch-clamp recordings in mouse spinal cord slices combined with selective pharmacological manipulation of NaV1.6, CaV1.3, KCNQ, Kv1.2, and HCN channels, we aimed to disentangle the relative contributions of inward and outward currents to nonlinear firing behaviors. We hypothesized that while PIC amplitude and hysteresis would increase with development, other conductances would play a decisive role in shaping recruitment–derecruitment asymmetry, providing a broader mechanistic basis for hysteresis than inward currents alone.

## Results

### PICs and recruitment-derecruitment hysteresis increase in fast motoneurons around weight bearing stages

Whole cell patch-clamp electrophysiology was deployed to study persistent inward currents (PICs) and firing hysteresis of lumbar motoneurons in transverse spinal cord slices (Figure 1A) obtained from mice during the second and third postnatal weeks. In these preparations, two motoneuron subtypes can be identified based on delayed and immediate firing profiles (Figure 1B). These two motoneuron subtypes display electrophysiological properties and molecular markers that are consistent with fast and slow motoneurons respectively (Leroy *et al*., 2014; Bos *et al*., 2018), with functional differences between these two subtypes emerging sometime during the second postnatal week (Sharples & Miles, 2021; Sharples *et al*., 2025). This period of time coincides with drastic changes in locomotor behaviour, with the emergence of hindlimb weightbearing toward the end of the second postnatal week (Altman & Sudarshan, 1975).

**Figure 1:**
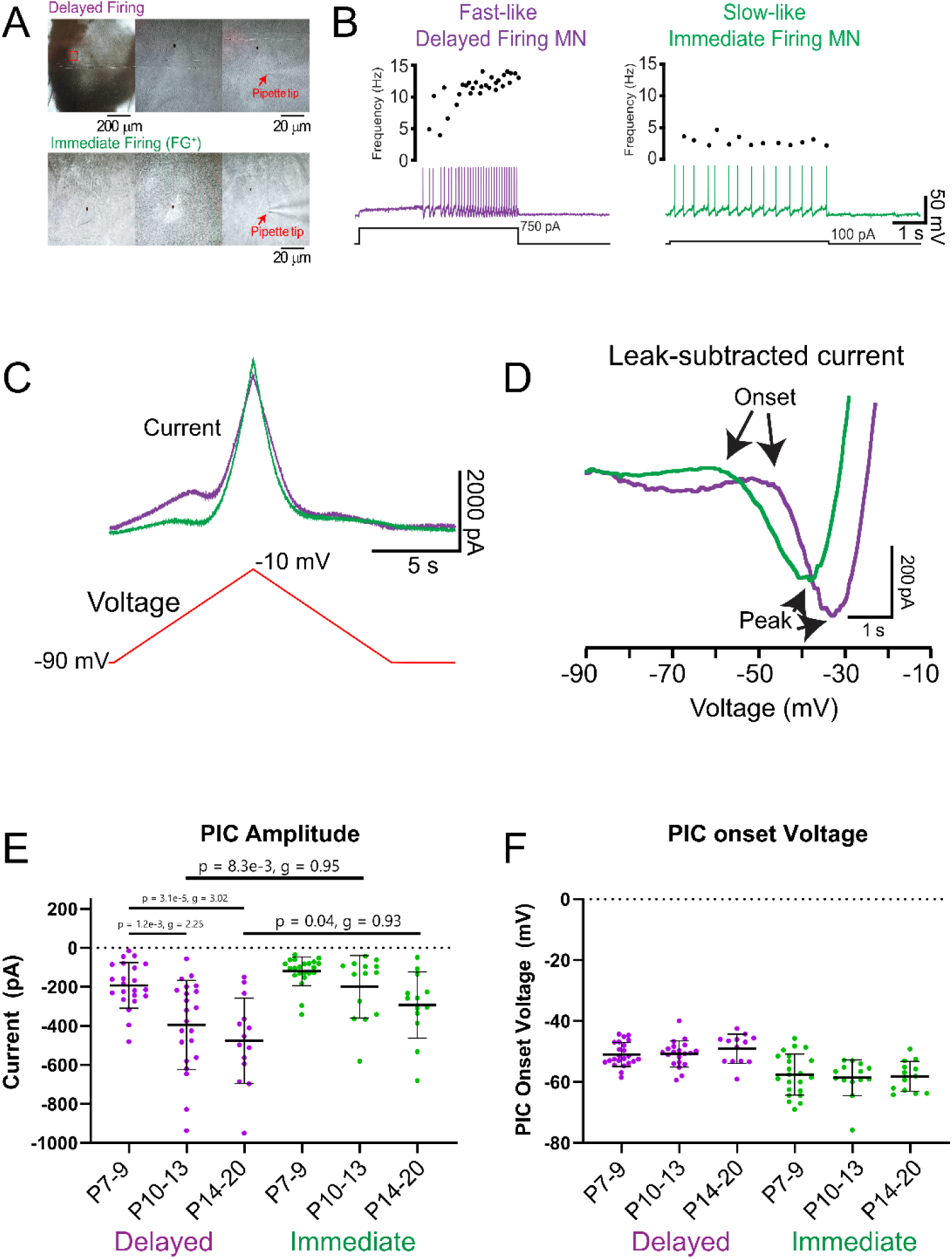
PICs increase in delayed firing motoneurons after the emergence of hindlimb weight bearing. (A) Motoneurons visualized under differential interference contrast were identified based on delayed (purple) and immediate (green) firing with long (5 second) depolarizing current steps applied near rheobase current (B). (C) Persistent inward currents (PICs) were measured in voltage clamp from delayed (purple) and immediate firing (green) motoneurons at pre (Delayed n = 22; immediate n = 22; P7-9) and post- (Delayed n = 22; immediate n = 14; P10-13) weight bearing stages and into the third postnatal week (Delayed n = 14; immediate n = 13; P 14-20) using a slow depolarizing voltage ramp (10mV/s; -90 to -10 mV). (F) PIC onset and amplitude were measured from leak-subtracted, low pass filtered (5Hz Bessel) traces. A-D, adapted from (Sharples & Miles, 2021). (G) PIC amplitude increased in delayed firing motoneurons from pre- to post-weight bearing stages and did not change in immediate firing motoneurons. (F) PIC onset voltage did not change in delayed or immediate firing motoneurons during the second and third postnatal week. Data are presented as mean ± SD with individual data points for each motoneuron displayed. Data were analyzed with a 2 factor ANOVA, with age and motoneuron type as factors. P values are reported when significant differences were detected. Hedge’s g values indicated small (0.2-0.49), medium (0.5-0.79) and large (>0.8) effect sizes.

Previous work has suggested that increases in PICs in motoneurons may contribute to the emergence of hindlimb weight-bearing, toward the end of the second postnatal week, by supporting sustained activation of the fatigue resistant slow twitch motor units that maintain tone in postural muscles of the hindlimb (Brocard *et al*., 1999; Vinay *et al*., 2000; Clarac *et al*., 2004; Jean-Xavier *et al*., 2018). To test this hypothesis, we reanalyzed a data set of 209 motoneurons from 81 mice from our previously published work (Sharples & Miles, 2021; Sharples *et al*., 2023), which measured PICs in voltage clamp during triangular voltage ramps (Figure 1C, D). In our previous work, all motoneurons studied during week 2 were pooled. In the current analysis, PICs were examined before and after the emergence of weight bearing during the early (P7-9) and late (P10-13) stages of the second and into the third (P14-20) postnatal weeks. In line with our hypothesis, PIC amplitude increased after weight bearing stages (P10-13) but did not continue to increase further into the third postnatal week (Figure 1E; Table 1a). However, in contrast to our hypothesis, and in line with previous work (Sharples & Miles, 2021; Harris-Warrick *et al*., 2023), PICs were larger in the ‘fast-like’ delayed-firing motoneurons compared to ‘slow-like’ immediate firing motoneurons (Table 2). This difference in PIC amplitude between delayed and immediate firing motoneurons emerged at the end of week 2 and was preserved into week 3 (Figure 1E). Interestingly, the majority of the PIC was activated below the spike threshold, however this proportion of PIC activated below spike threshold did not differ between motoneuron subtypes or change across weeks 2 and 3 (Table 2; Table 1b). PIC onset voltage was more depolarized in delayed compared to immediate firing motoneurons but did not change across these three developmental time points (Figure 1F; Table 2; Table1c).

**Table 1:**
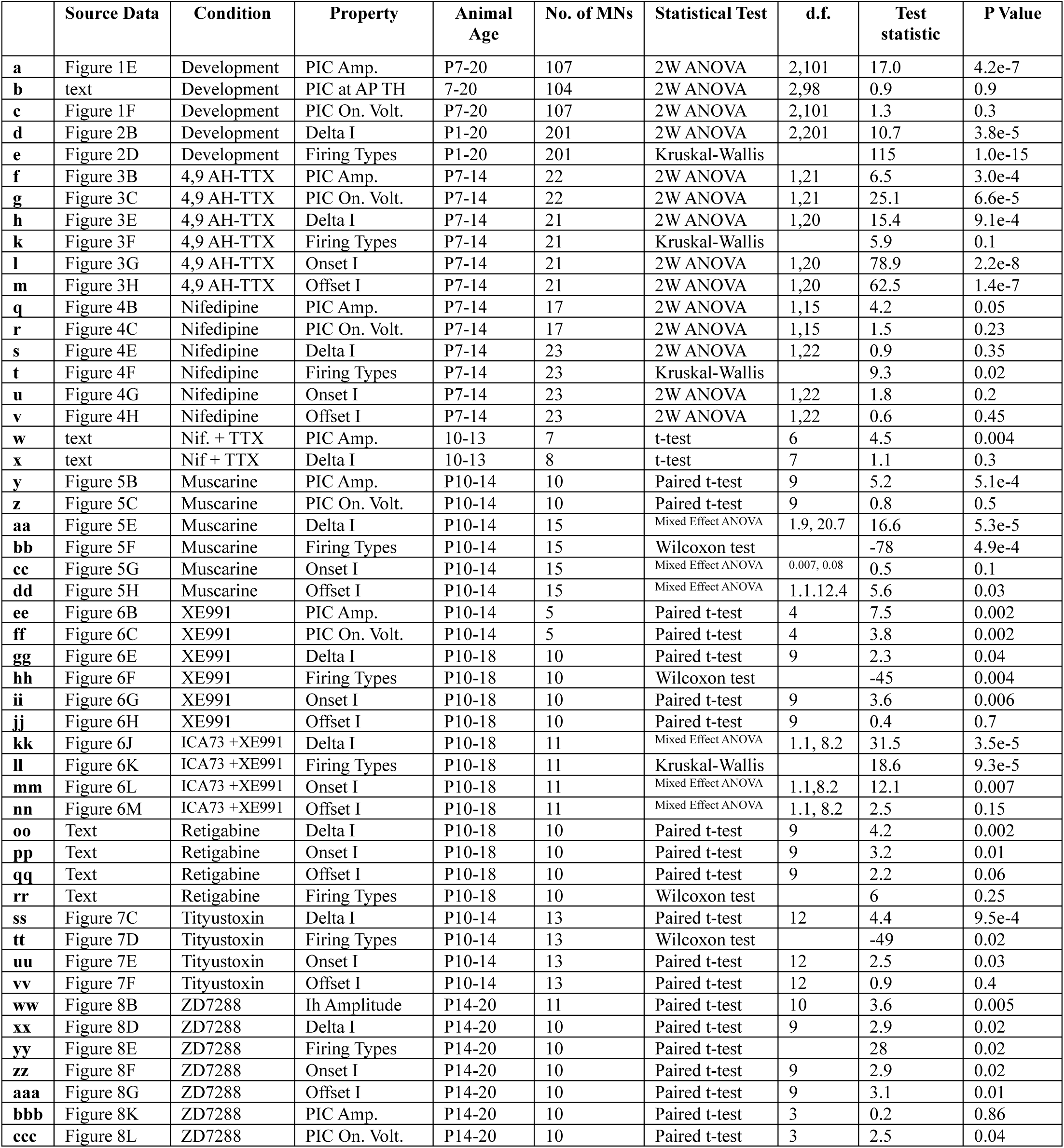
Statistical summary table. Results from statistical tests performed in this study are annotated through the manuscript and are indicated in the first column of the table.

**Table 2:**
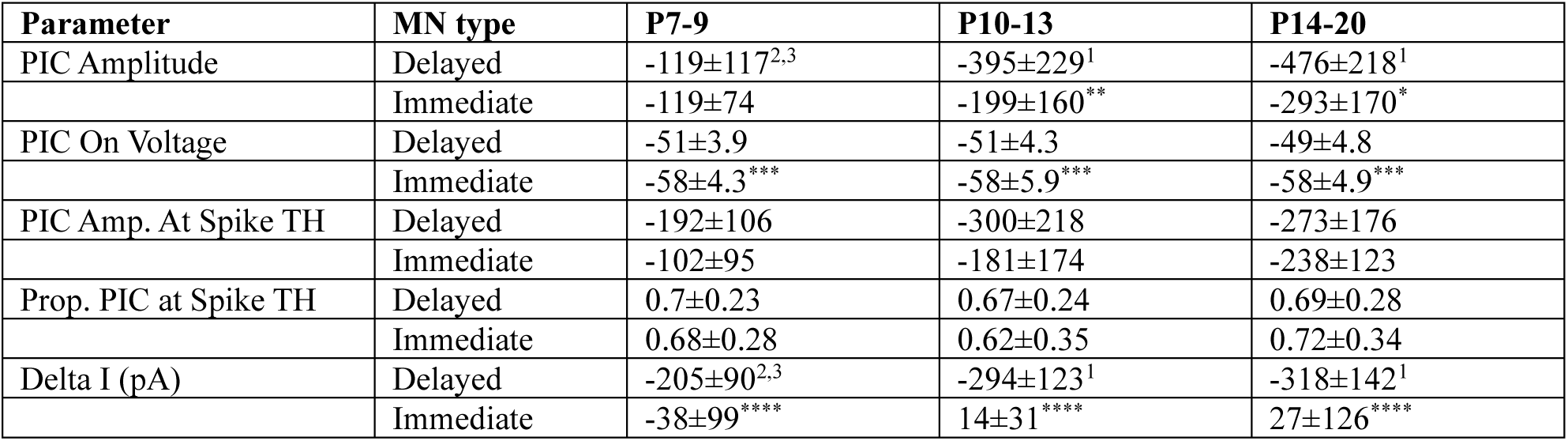
Developmental analysis of PIC and recruitment-derecruitment hysteresis. Data are presented as mean ± SD. Superscript numbers indicate significant differences within motoneuron subtypes between P7-9 (1), P10-13 (2), or P14-20 (3). Asterisks denote significant differences between motoneuron subtypes within each developmental stage with *p<0.05, **p<0.01, ***p<0.001, ****p<0.00001.

| Parameter | MN type | P7-9 | P10-13 | P14-20 |
| --- | --- | --- | --- | --- |
| PIC Amplitude | Delayed | -119±117<sup>2,3</sup> | -395±229<sup>1</sup> | -476±218<sup>1</sup> |
| | Immediate | -119±74 | -199±160** | -293±170* |
| PIC On Voltage | Delayed | -51±3.9 | -51±4.3 | -49±4.8 |
| | Immediate | -58±4.3*** | -58±5.9*** | -58±4.9*** |
| PIC Amp. At Spike TH | Delayed | -192±106 | -300±218 | -273±176 |
| | Immediate | -102±95 | -181±174 | -238±123 |
| Prop. PIC at Spike TH | Delayed | 0.7±0.23 | 0.67±0.24 | 0.69±0.28 |
| | Immediate | 0.68±0.28 | 0.62±0.35 | 0.72±0.34 |
| Delta I (pA) | Delayed | -205±90<sup>2,3</sup> | -294±123<sup>1</sup> | -318±142<sup>1</sup> |
| | Immediate | -38±99**** | 14±31**** | 27±126**** |

In addition to measuring PICs in voltage clamp, we also assessed recruitment-derecruitment and firing rate hysteresis in current clamp during slow triangular currents ramps (Figure 2A). This is an approach that is often used to estimate the influence of PICs on motoneuron firing, with negative values reflective of firing persisting at lower levels of current on the descending limb of the ramp compared to the ascending limb of the ramp (Ioff<Ion) (Li & Bennett, 2003; Li *et al*., 2004; Harvey *et al*., 2006b; Quinlan *et al*., 2011; Durand *et al*., 2015; Steele *et al*., 2020; Sharples & Miles, 2021; Harris-Warrick *et al*., 2023). In line with measurements of PICs made in voltage clamp, recruitment-derecruitment hysteresis (delta I) became more negative (increased) in delayed firing motoneurons but did not change in immediate firing motoneurons across the second and third weeks of postnatal development (Figure 2B; Table 2; Table 1d). Delta I plateaued at the end of the second postnatal week, with no further change into week 3 (Figure 2B; Table 1d).

**Figure 2:**
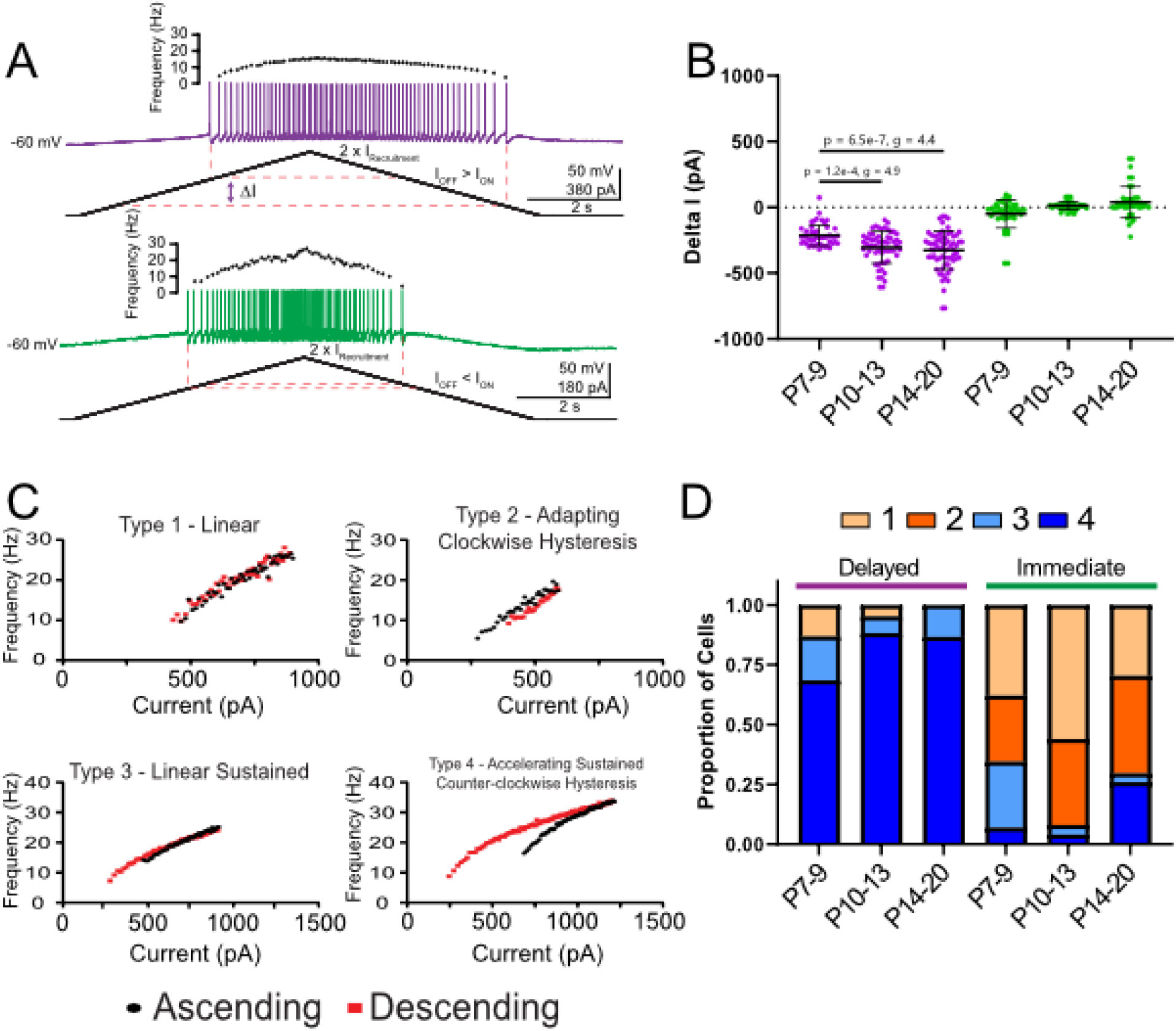
Recruitment-derecruitment and firing hysteresis mature in parallel to PICs. (A) Representative traces of the membrane potential and repetitive firing from a delayed (purple) and immediate firing (green) motoneuron during a triangular depolarizing current ramp up to an intensity of 2 x their respective rheobase currents. (B) Recruitment-derecruitment hysteresis, measured as Delta I, represents the difference between current at firing offset on the descending limb of the ramp and the current at firing onset on the ascending limb of the ramp, increased in delayed firing motoneurons at pre- and post-weight bearing stages but did not change in immediate firing motoneurons. (C) Frequency current plots with firing rates derived from ascending (black) and descending (red) limbs of triangular current ramps. Four patterns of firing rate hysteresis can be identified as described by Li and Bennett, 2006; with types 3 and 4 suggestive of PIC actions. A&C, adapted from (Sharples & Miles, 2021). (D) Relative proportion of firing hysteresis types in delayed and immediate firing motoneurons across the second and third postnatal week. Data are presented as mean ± SD with individual data points for each motoneuron displayed. Data in B were analyzed with a 2 factor ANOVA, with age and motoneuron type as factors. P values are reported when significant differences were detected. Hedge’s g values indicated small (0.2-0.49), medium (0.5-0.79) and large (>0.8) effect sizes. (Delayed firing MNs: P7-9 n = 31, P10-13 n = 43, P14-20 n = 52; Immediate firing MNs: P7-9 n=29, P10-13 n = 26, P14-20 n = 27).

Motoneurons also present with 4 classes of firing rate hysteresis during triangular current ramps, with type 1; linear, type 2; adapting/clockwise hysteresis, type 3; sustained, and type 4; counterclockwise hysteresis (Figure 2C). Of these classes, types 3 and 4 have been proposed to be mediated by a PIC (Bennett *et al*., 2001; Durand *et al*., 2015; Sharples & Miles, 2021). Overall, these firing types did not significantly change within either subtype over the first three postnatal weeks, however significant differences were found between delayed and immediate firing motoneurons at all time points (Figure 2D; Table 1e).

### Nav1.6 and L-type calcium channels contribute to PICs but do not underlie hysteresis in fast motoneurons

Having identified a time course that PIC, recruitment-derecruitment, and firing hysteresis mature during postnatal development, we next set out to determine which PIC-conducting ion channels contribute to the increase in PICs and recruitment-derecruitment hysteresis in delayed firing motoneurons around weight bearing stages (P7-9 vs. P10-13).

Repetitive firing during slow depolarizing current injection is critically dependent on the availability of persistent sodium channels (INaP) and can be blocked with riluzole (Miles *et al*., 2005; Kuo *et al*., 2006). Because persistent sodium current is a major contributor to PIC amplitude, reducing this current would be expected to decrease PIC amplitude and consequently reduce delta I, reflecting a diminished contribution of PICs to firing hysteresis. Consistent with our previous findings (Sharples & Miles, 2021) and those of others (Drouillas *et al*., 2023), blocking NaV1.6 channels produced a 34 ± 40% reduction in PIC amplitude at P7–9 and a 39 ± 35% reduction at P10–13, with no difference in the magnitude of this reduction between developmental stages (Figure 3A,B; Table 3; Table 1f). Blocking NaV1.6 channels also depolarized the onset voltage of the PIC at both early and late stages of week 2, again with no difference in the magnitude of this shift between stages (Figure 3C; Table 1g). Despite the reduction in PIC amplitude, however, delta I increased following NaV1.6 blockade (Figure 3E; Table 1h), contrary to the prediction that reducing PIC amplitude should decrease delta I. This increase in delta I was observed at both early and late stages of week 2 and occurred without a change in the proportions of Type 1-4 firing hysteresis (Figure 3F; Table 1k). The increase in delta I was paralleled by increases in both recruitment and derecruitment currents (Figure 3G, H; Table 1l,m). Together, these data indicate that NaV1.6 channels contribute to the amplitude and shape the onset voltage of the underlying PIC. However, changes in NaV1.6 function do not account for the developmental increase in PIC amplitude during the second postnatal week and appear to contribute minimally to recruitment–derecruitment dynamics and firing hysteresis.

**Figure 3:**
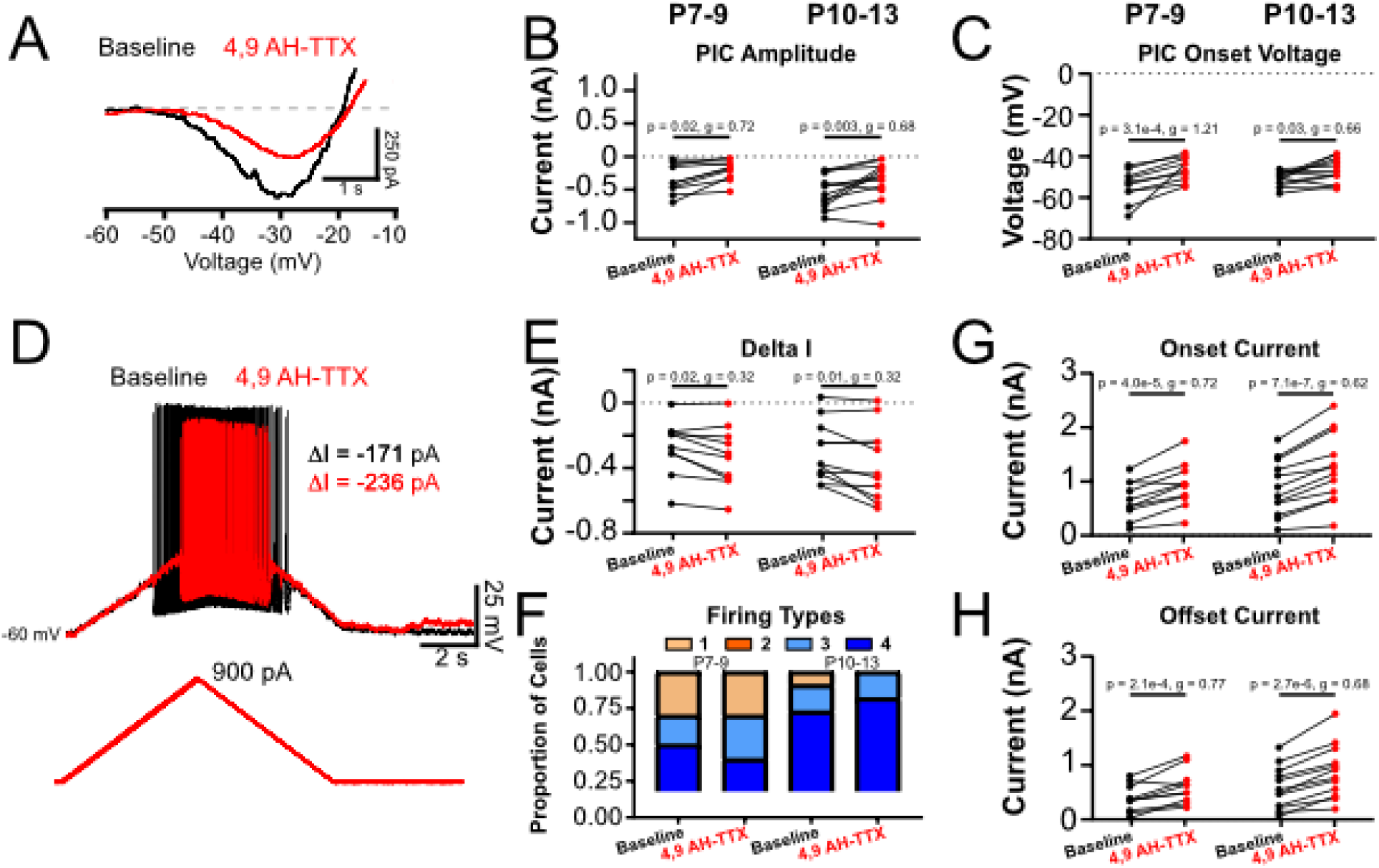
Nav 1.6 channels contribute to PIC but not firing hysteresis. (A) Representative traces of a leak-subtracted and filtered PIC measured in voltage clamp before (black) and after (red) application of the NaV1.6 blocker 4,9 Anhydro-tetrodotoxin (4,9 AH-TTX; 200 nM). (B) 4,9 AH-TTX decreased PIC amplitude and depolarized PIC onset voltage (C) at both pre- (n=10; P7-9) and post- (n=12; P10-13) weight bearing stages. (D) Representative traces of the membrane potential and repetitive firing during a triangular depolarizing current ramp before (black) and after (red) application of 4,9 AH-TTX. (E) 4,9 AH-TTX increased recruitment-derecruitment hysteresis (Delta I) but did not alter the proportion of firing types (F) at pre- and post-weight bearing stages. 4,9 AH-TTX increased the current at firing onset on the ascending limb of the ramp (G) and increased current at firing offset on descending limb of the ramp (H) at both pre-and post-weight bearing stages. Data are presented as individual data points and were analyzed using a 2 factor ANOVA with drug and stage as factors. P values are reported when significant differences were detected. Hedge’s g values indicated small (0.2-0.49), medium (0.5-0.79) and large (>0.8) effect sizes.

**Table 3:**
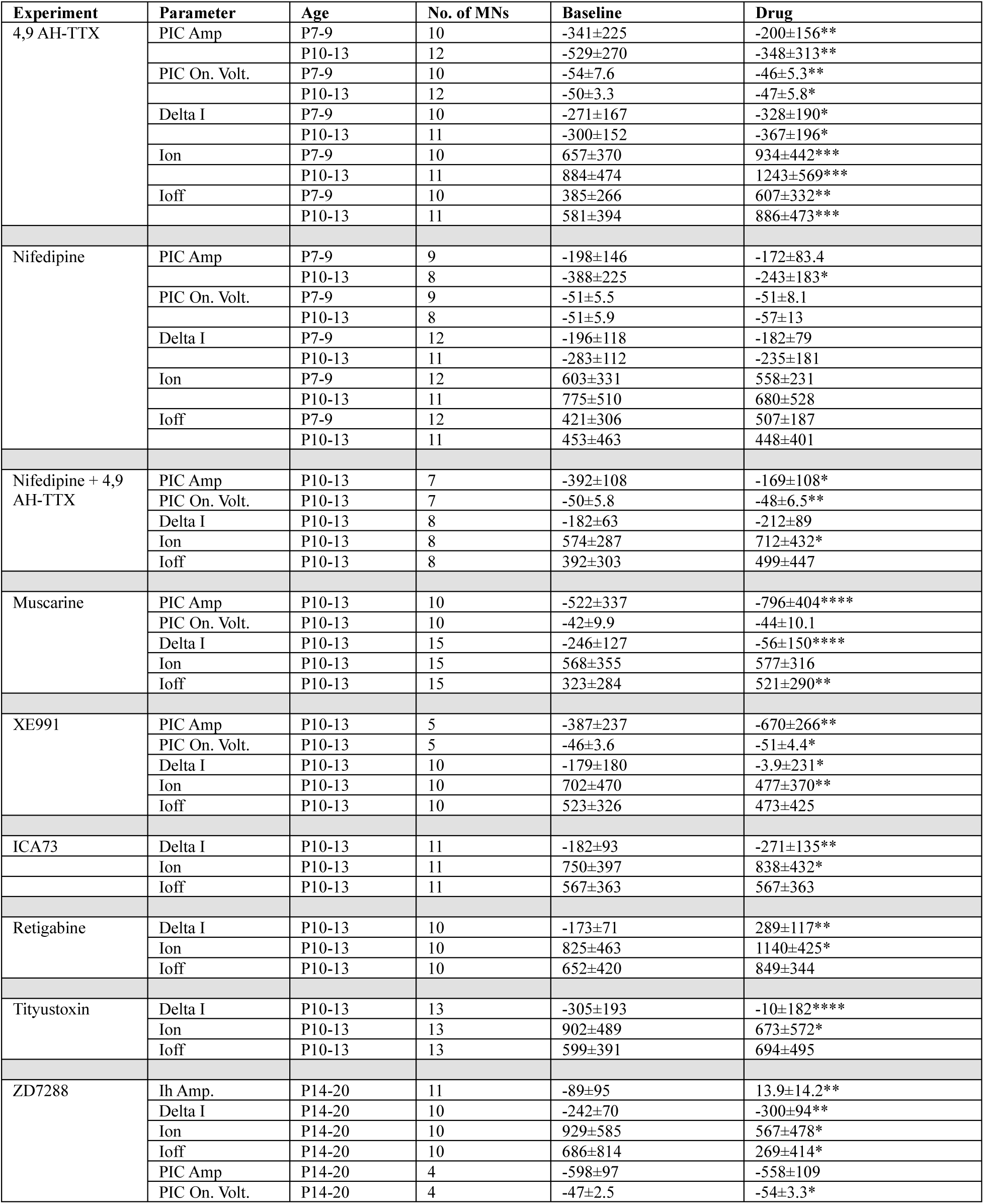
Pharmacological manipulation of PIC measured in voltage clamp and recruitment-derecruitment hysteresis measured in current clamp. Data are presented as mean ± SD. Asterisks denote significant differences between drug and baseline conditions with *p<0.05, **p<0.01, ***p<0.001, ****p<0.00001.

We next tested the contribution of L-type calcium channels to the PIC measured during the second postnatal week to determine if changes in their contribution can account for the increase in PIC amplitude and recruitment-derecruitment hysteresis during the second postnatal week. Blocking L-type calcium channels with nifedipine did not alter PIC amplitude at P7-9 (3±20%) but produced a 34±26% reduction in the amplitude of the PIC at P10-13 (Figure 4A, B; Table 3; Table 1q). Consistent with previous reports (Sharples & Miles, 2021), nifedipine did not change the onset voltage of the PIC at either stage (Figure 4C; Table 1r). Despite the reduction in PIC amplitude, nifedipine did not significantly change delta I (Figure 4D, E; Table 1s), proportions of firing hysteresis (Figure 4F; Table 1t), onset current (Figure 4G; Table 1u), or offset current (Figure 4H; Table 1v) at either time point. Together these data suggest that L-type calcium channels contribute in part to PIC amplitude and an increase in their contribution toward the end of the second postnatal week may account for the increase in PIC amplitude that we observed. However, in contrast to what we expected, L-type calcium channels are not a key contributor to recruitment-derecruitment or firing hysteresis.

**Figure 4:**
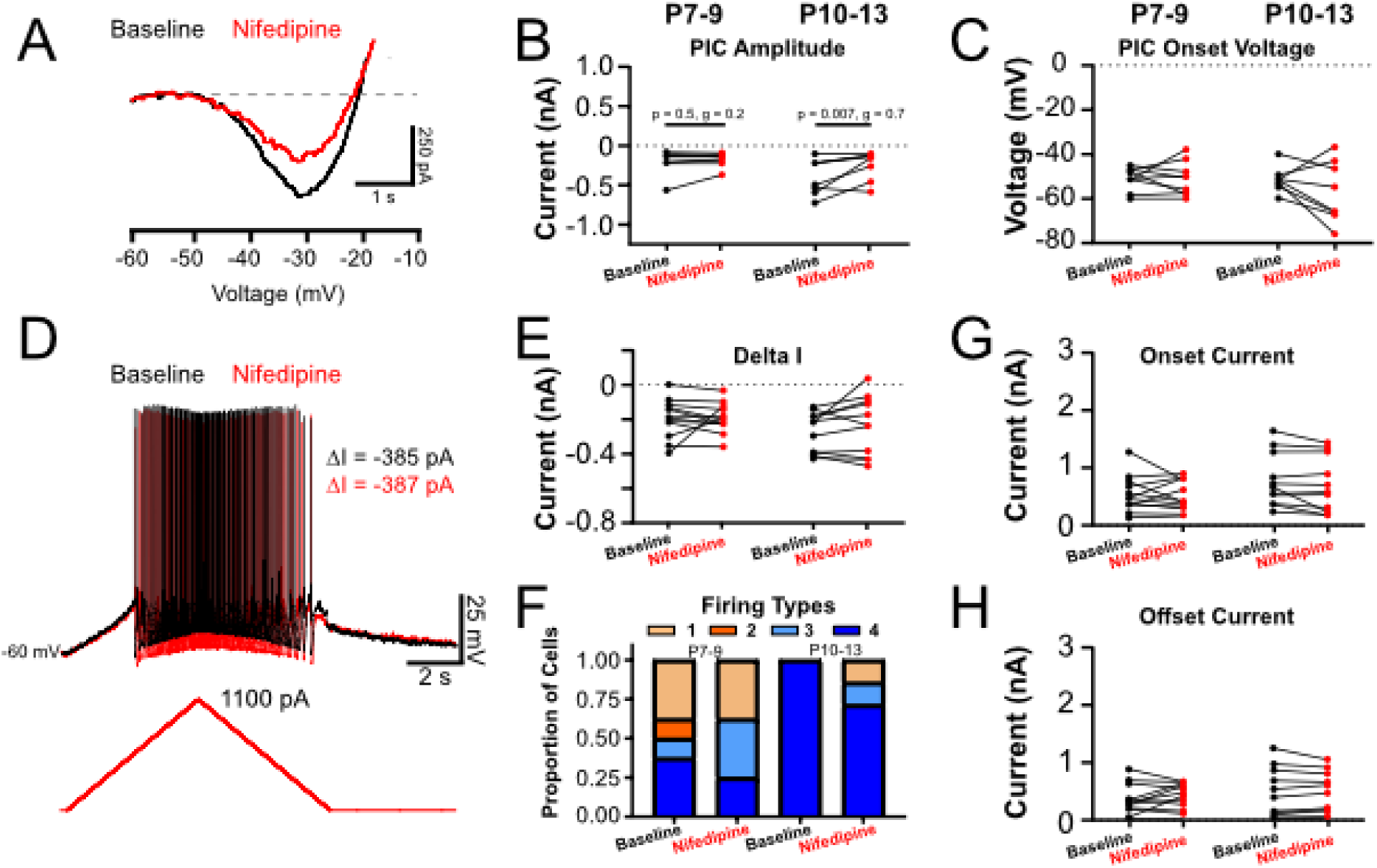
L-type calcium channels contribute to PIC after the emergence of weight bearing but do not contribute to firing hysteresis. (A) Representative traces of a leak-subtracted and filtered PIC measured in voltage clamp before (black) and after (red) application of the L-type calcium channel blocker Nifedipine (20 µM). (B) Nifedipine decreased PIC amplitude at post weight-bearing stages but did not alter PIC onset voltage (C). (D) Representative traces of the membrane potential and repetitive firing during a triangular depolarizing current ramp before (black) and after (red) application of Nifedipine. (E) Nifedipine did not affect recruitment-derecruitment hysteresis (Delta I) or alter the proportion of firing types (F) at pre- (n= 12; P7-9) and post-weight bearing stages (n= 11; P10-13). Nifedipine did not affect current at firing onset on the ascending limb of the ramp (G) or current at firing offset on descending limb of the ramp (H) at both pre- or post-weight bearing stages. Data are presented as individual data points and were analyzed using a 2 factor ANOVA with drug and stage as factors. P values are reported when significant differences were detected. Hedge’s g values indicated small (0.2-0.49), medium (0.5-0.79) and large (>0.8) effect sizes.

We next considered the possibility that blocking NaV1.6 or L-type calcium channels individually did not produce a sufficient reduction in PIC amplitude to alter firing hysteresis. If delta I is strongly dependent on PIC magnitude, then a larger reduction in PIC amplitude should be expected to decrease delta I. To test this possibility, we simultaneously blocked NaV1.6 and L-type calcium channels using 4,9-anhydro-tetrodotoxin and nifedipine. Although co-application of these antagonists (n = 8 MNs; P10–13) produced a substantial 58 ± 7.7% reduction in PIC amplitude (Table 3; Table 1w), this manipulation did not significantly alter delta I or firing hysteresis in the expected direction (Table 3; Table 1x). These findings further indicate that the magnitude of the PIC alone is not a primary determinant of recruitment–derecruitment hysteresis in developing motoneurons.

### Paradoxical modulation of PIC amplitude and firing hysteresis in fast motoneurons

Previous work has suggested that the influence of PICs on motoneuron firing is critically dependent on the presence of endogenous sources of neuromodulation (Hounsgaard *et al*., 1984, 1988; Conway *et al*., 1988). It has also been shown that multiple neuromodulators can increase PIC amplitude in hypoglossal motoneurons (Revill *et al*., 2019). Here we tested the hypothesis that increasing PIC amplitude by activating muscarinic acetylcholine receptors would produce a shift toward sustained firing hysteresis, as we found during development. In line with previous reports (Revill and Funk, 2022), application of muscarine produced a 174±82% increase in PIC amplitude (Figure 5A, B; Table 3; Table 1y) but did not alter the onset voltage of the PIC (Figure 5A, C; Table 1z). This increase in PIC amplitude produced by muscarine is equivalent to the mean increase that we observed between the early and late phase of the second postnatal week. However, in contrast to what would be expected, muscarine decreased delta I (Figure 5D, E; Table 1aa) and produced a general shift from types 3-4 to types 1-2 firing hysteresis (Figure 5F; Table 1bb)). The reduction in delta I occurred without a change in onset current (Figure 5G; Table 1cc) and was instead attributable to a selective increase in the offset current in the presence of muscarine (Figure 5H; Table 1dd).

**Figure 5:**
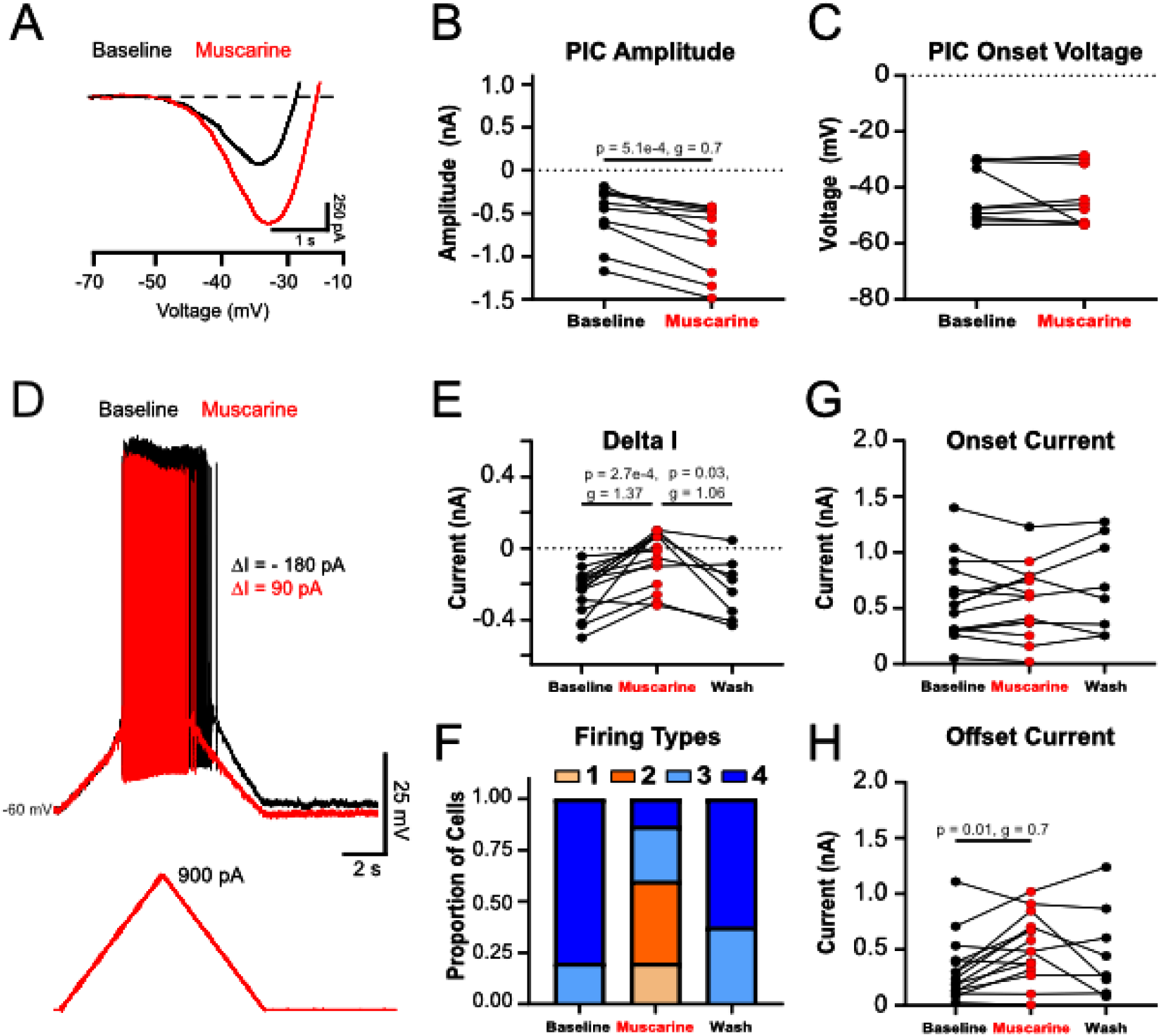
Muscarine increases PIC but reduces firing hysteresis. (A) Representative traces of a leak-subtracted and filtered PIC measured in voltage clamp before (black) and after (red) application of the Muscarine (10 µM; n = 10; P7-13). (B) Muscarine increased PIC amplitude but did not alter PIC onset voltage (C). (D) Representative traces of the membrane potential and repetitive firing during a triangular depolarizing current ramp before (black) and after (red) application of muscarine. (E) Muscarine decreased recruitment-derecruitment hysteresis (Delta I; n = 15) and produced a shift in firing types from types 3 and 4 to types 1-4. These effects were reversed following a wash with regular aCSF. Muscarine did not affect current at firing onset on the ascending limb of the ramp (G) but increased the current of firing offset on descending limb of the ramp (H). Data are presented as individual data points and were analyzed using a paired t-test (B & C) or repeated measures ANOVA (E - H). P values are reported when significant differences were detected. Hedge’s g values indicated small (0.2-0.49), medium (0.5-0.79) and large (>0.8) effect sizes.

### Firing hysteresis in fast motoneurons is modulated by potassium channels

Although muscarine increases PIC amplitude, it is also known to modulate multiple voltage-sensitive ion channels. One current of particular interest is the M-type potassium current, conducted by slow-activating, non-inactivating KCNQ channels that are largely active in the subthreshold voltage range (Alaburda *et al*., 2002; Ghezzi *et al*., 2017, 2018; Verneuil *et al*., 2020; Sharples *et al*., 2023). Because these channels a persistent outward current, it can oppose the actions of PICs (Verneuil *et al*., 2020; Singh *et al*., 2025; Gaudreau & Bui, 2026) and was originally identified based on its inhibition by muscarinic receptor activation (Brown & Adams, 1980). In addition, our previous work demonstrated large M-currents in delayed firing motoneurons and a selective role for this conductance in controlling recruitment (Sharples *et al*., 2023).

Based on these observations, we hypothesized that M-currents shape firing hysteresis by counteracting PICs and by providing differential control of recruitment and derecruitment currents (Verneuil *et al*., 2020; Sharples *et al*., 2023). Under this framework, reducing M-current would be expected to enhance PIC amplitude and potentially increase delta I by strengthening inward current amplification during firing. Consistent with the predicted effect on PICs, blocking M-currents with XE991 (10 μM) produced a 188 ± 32% increase in PIC amplitude (Figure 6A, B; Table 3; Table 1ee) and hyperpolarized PIC onset voltage by 4.9 ± 2.8 mV (Figure 6C; Table 1ff). The magnitude of this increase in PIC amplitude was comparable to that observed developmentally and following muscarinic activation. However, despite the substantial increase in PIC amplitude, blocking KCNQ channels reduced delta I (Figure 6D, E; Table 1gg) and shifted firing behavior from Types 3–4 toward Types 1–3 patterns (Figure 6F; Table 1hh), contrary to the expectation that larger PICs should increase firing hysteresis. Importantly, the mechanism underlying this reduction in delta I differed from that observed with muscarine. Following KCNQ channel blockade, the reduction in delta I resulted from a decrease in the recruitment current (Figure 6G; Table 1ii), with no change in the derecruitment current (Figure 6H; Table 1jj). Thus, whereas muscarine reduced delta I by increasing the offset current, inhibition of KCNQ channels reduced delta I by lowering the onset current on the ascending limb of the current ramp.

**Figure 6:**
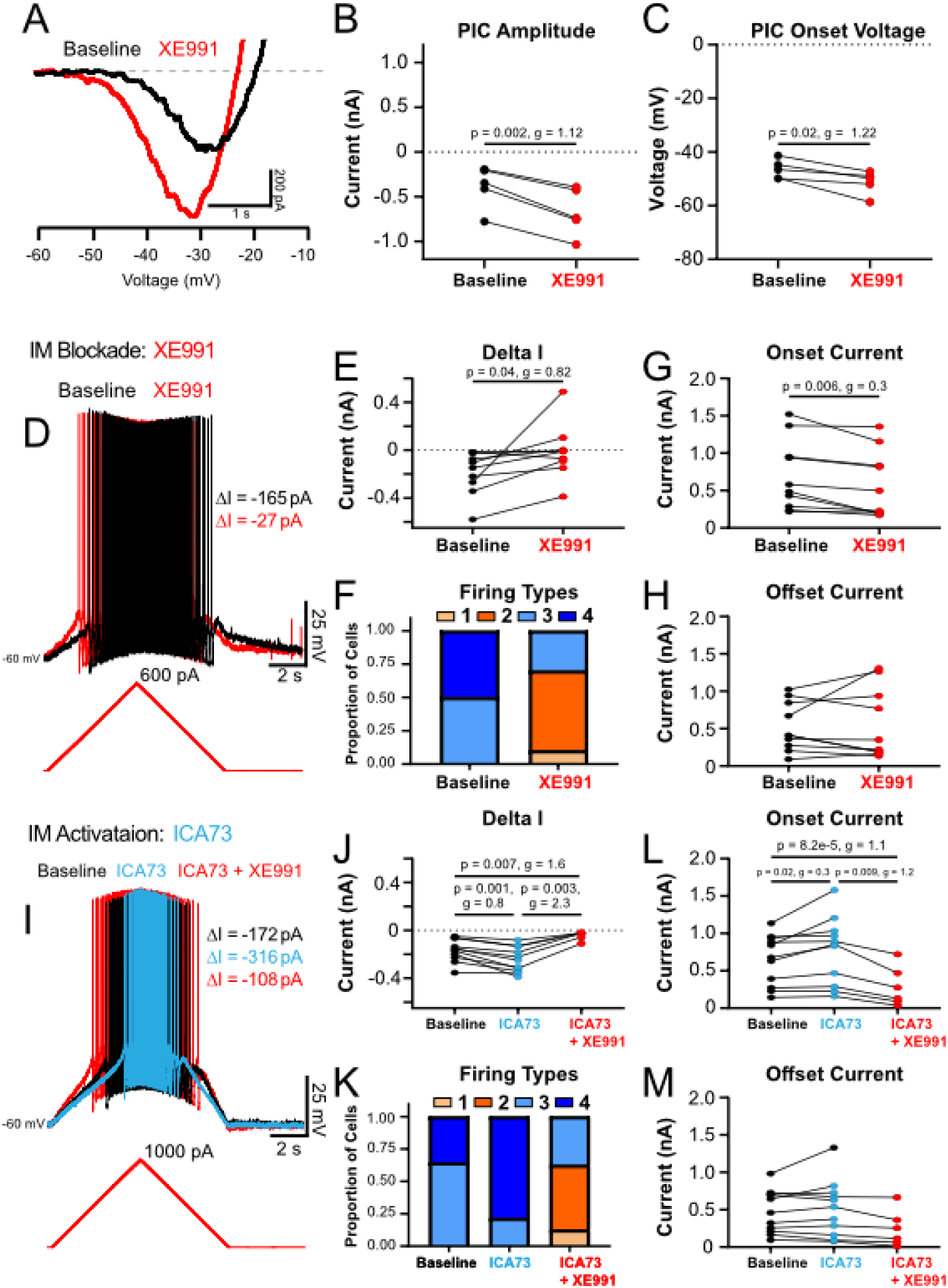
KCNQ channels attenuate PIC and promote firing hysteresis. (A) Representative traces of a leak-subtracted and filtered PIC measured in voltage clamp before (black) and after (red) application of the KCNQ channel blocker XE991 (10 µM; n = 5; P10-13). (B) XE991 increased PIC amplitude and hyperpolarized PIC onset voltage (C). (D) Representative traces of the membrane potential and repetitive firing during a triangular depolarizing current ramp before (black) and after (red) application of XE991. (E) XE991 decreased recruitment-derecruitment hysteresis (Delta I; n = 10) and produced a shift in firing types from types 3 and 4 to types 1-3 (F). XE991 decreased the current at firing onset on the ascending limb of the ramp (G) but did not alter the current of firing offset on the descending limb of the ramp (H). (I) Representative traces of the membrane potential and repetitive firing during a triangular depolarizing current ramp before (black) and after (blue) application of the KCNQ channel activator ICA73 (n=11; 10 µM) and subsequent application of XE991 (red). (J) ICA73 increased recruitment-derecruitment hysteresis (Delta I) and produced a shift in firing types from types 3 and 4 to types 1-3 (K). ICA73 increased the current at firing onset on the ascending limb of the ramp (L) but did not alter the current of firing offset on the descending limb of the ramp (M). These effects were reversed with subsequent application of the KCNQ channel blocker, XE991 (n = 6). Data are presented as individual data points and were analyzed using a paired t-test (B & C) or repeated measures ANOVA (E - H; J - M). P values are reported when significant differences were detected. Hedge’s g values indicated small (0.2-0.49), medium (0.5-0.79) and large (>0.8) effect sizes.

Conversely, activation of KCNQ channels with ICA73 increased delta I (Figure 6I, J; Table 3; Table 1kk) and produced a general shift toward Type 4 firing hysteresis (Figure 6K; Table 1ll). This increase in delta I resulted from an increase in recruitment current (Figure 6L; Table 1mm) with no change in derecruitment current (Figure 6M; Table 1nn). Similar effects were observed with a second KCNQ channel activator, retigabine (Table 3; Table 1oo-rr). Moreover, these effects were reversed by subsequent application of the KCNQ blocker XE991, which reduced delta I to levels significantly below baseline and shifted firing behavior toward Types 1–2 patterns, consistent with blockade of KCNQ channels alone (Figure 6J–M; Table 1gg-jj). Together, these findings indicate that KCNQ channels conduct an outward current that not only opposes PICs but also plays a critical role in shaping recruitment–derecruitment hysteresis. Rather than regulating derecruitment, KCNQ channels primarily influence firing hysteresis by controlling the recruitment current on the ascending limb of triangular current ramps.

Given that types 3-4 firing hysteresis and negative delta I values are a prominent feature of fast-type delayed firing motoneurons (Figure 2), with delayed firing being produced by rapid activation and slow inactivation of Kv1.2 channels (Bos *et al*., 2018; Harris-Warrick *et al*., 2024), we next tested the hypothesis that slow inactivation of Kv1.2 channels might contribute to sustained firing hysteresis by allowing motoneurons to fire for longer on the descending aspect triangular current ramps. Blockade of Kv1.2 channels with tityustoxin (TsTx; 800 nM) blocked the slow depolarization of the membrane potential in response to a long (5s) depolarizing current step applied near rheobase (Figure 7A). Interestingly, and in line with our hypothesis, blocking Kv1.2 channels decreased delta I (Figure 7B, C; Table 3; Table 1ss) and produced a shift from Types 3 and 4 to Types 1-3 firing hysteresis (Figure 7D; Table 1tt). However, in contrast to what we hypothesized, the reduction of delta I was due to a significant decrease in the onset current (Figure 7E; Table 1uu) and no change in the offset current (Figure 7F; Table 1vv). These data suggest that the rapid activation of Kv1.2 channels may influence firing hysteresis by shaping fast motoneuron recruitment, however slow inactivation of Kv1.2 channels may play less of a role in supporting sustained firing on the descending limb of the ramp.

**Figure 7:**
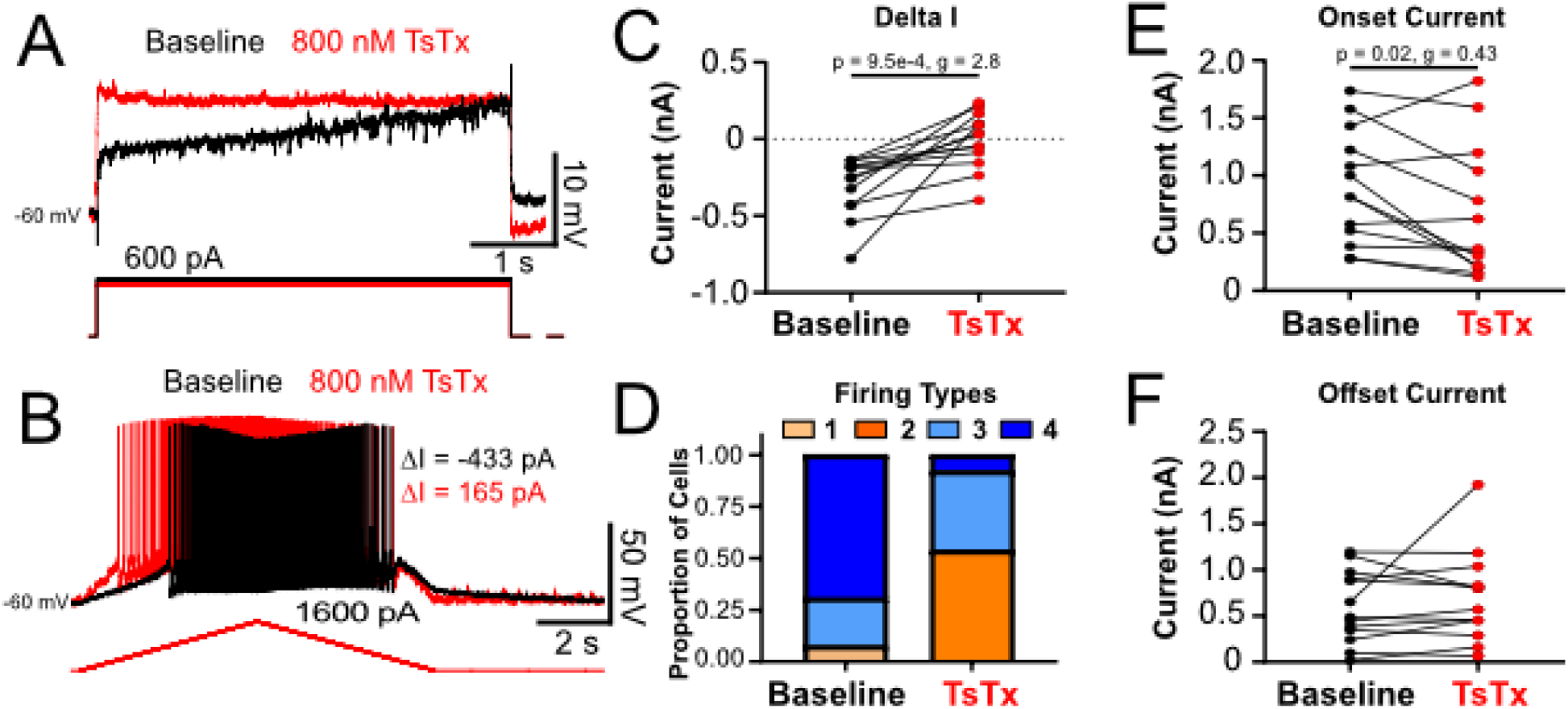
Kv1.2 channels shape firing hysteresis. (A) Representative traces of the membrane potential during a long (5 s) depolarizing current step applied just below rheobase before (black) and after (red) application of the Kv1.2 channel blocker Tityustoxin (800 nM; TsTx; n = 13; P7-13). (B) Representative traces of the membrane potential and repetitive firing during a triangular depolarizing current ramp before (black) and after (red) application of TsTx. (C) TsTx decreased recruitment-derecruitment hysteresis (delta I) and produced a shift in firing types from types 1, 3 and 4 to types 2-4. TsTx decreased the current at firing onset on the ascending limb of the ramp (G) but did not affect the current of firing offset on descending limb of the ramp (H). Data are presented as individual data points and were analyzed using a paired t-test. P values are reported when significant differences were detected. Hedge’s g values indicated small (0.2-0.49), medium (0.5-0.79) and large (>0.8) effect sizes.

### HCN channels modulate firing hysteresis and prevent bistable firing in fast motoneurons

In the preceding experiments, we found that manipulating PIC amplitude alone did not reliably predict changes in recruitment–derecruitment hysteresis. Both NaV1.6 and KCNQ channel manipulations substantially altered PIC amplitude yet produced changes in delta I that could not be explained solely by the magnitude of the PIC. These findings suggested that additional conductances that shape the membrane operating range or input conductance may play an important role in determining whether PICs generate firing hysteresis.

In a final set of experiments, we therefore assessed the contribution of hyperpolarization-activated cyclic nucleotide-gated (HCN) channels to recruitment–derecruitment and firing rate hysteresis in delayed-firing fast motoneurons. Our previous work demonstrated that by the third postnatal week (P14–20) the activation voltage of HCN channels shifts in the depolarizing direction, resulting in an h-current that is active at resting membrane potential. This resting h-current acts as a depolarizing shunt conductance that delays recruitment in response to depolarizing input (Sharples & Miles, 2021). Because HCN channels deactivate slowly during depolarization and reactivate during hyperpolarization, we hypothesized that this current could limit the ability of PICs to generate bistability and thereby suppress recruitment–derecruitment hysteresis during triangular current ramps. Consistent with this hypothesis, pharmacological blockade of HCN channels with ZD7288 (Figure 8A, B; Table 3; Table 1ww) significantly increased delta I (Figure 8C, D; Table 1xx). Notably, following HCN channel blockade, 48% of motoneurons displayed self-sustained firing— a feature rarely observed in delayed- or immediate-firing motoneurons during the first three postnatal weeks (Sharples & Miles, 2021). This enhancement in sustained activity was reflected by a shift in firing hysteresis from type 4 at baseline to the emergence of a fifth firing pattern characterized by persistent self-sustained firing (Figure 8E; Table 1yy). The increase in delta I was accompanied by significant reductions in both recruitment (Figure 8F; Table 1zz) and derecruitment (Figure 8G; Table 1aaa) currents during triangular current ramps. Moreover, after termination of the ramp (Figure 8H) or a brief depolarizing current step (Figure 8I), sustained discharge required hyperpolarizing current injection to terminate firing, consistent with the emergence of bistable firing behavior following HCN channel blockade. To determine whether this enhanced bistability resulted from changes in the underlying PIC, we measured PIC amplitude in voltage clamp in a subset of motoneurons (n = 4). Because PICs are believed to be a key contributor to self-sustained firing, we predicted removing the shunt current by blocking HCN channels might increase PIC amplitude. However, in contrast to this prediction, HCN channel blockade produced no significant change in PIC amplitude (−10 ± 40%, Figure 8J, K; Table 1bbb). Instead, the onset voltage of the PIC became significantly hyperpolarized (−6.4 ± 5.1 mV, Figure 8L; Table 1ccc) in all cells examined. Together, these findings suggest that HCN channels oppose the generation of self-sustained firing in fast motoneurons at the third postnatal week, likely by acting as a resting shunt conductance that stabilizes membrane potential and limits the voltage range over which PICs can generate bistability.

**Figure 8:**
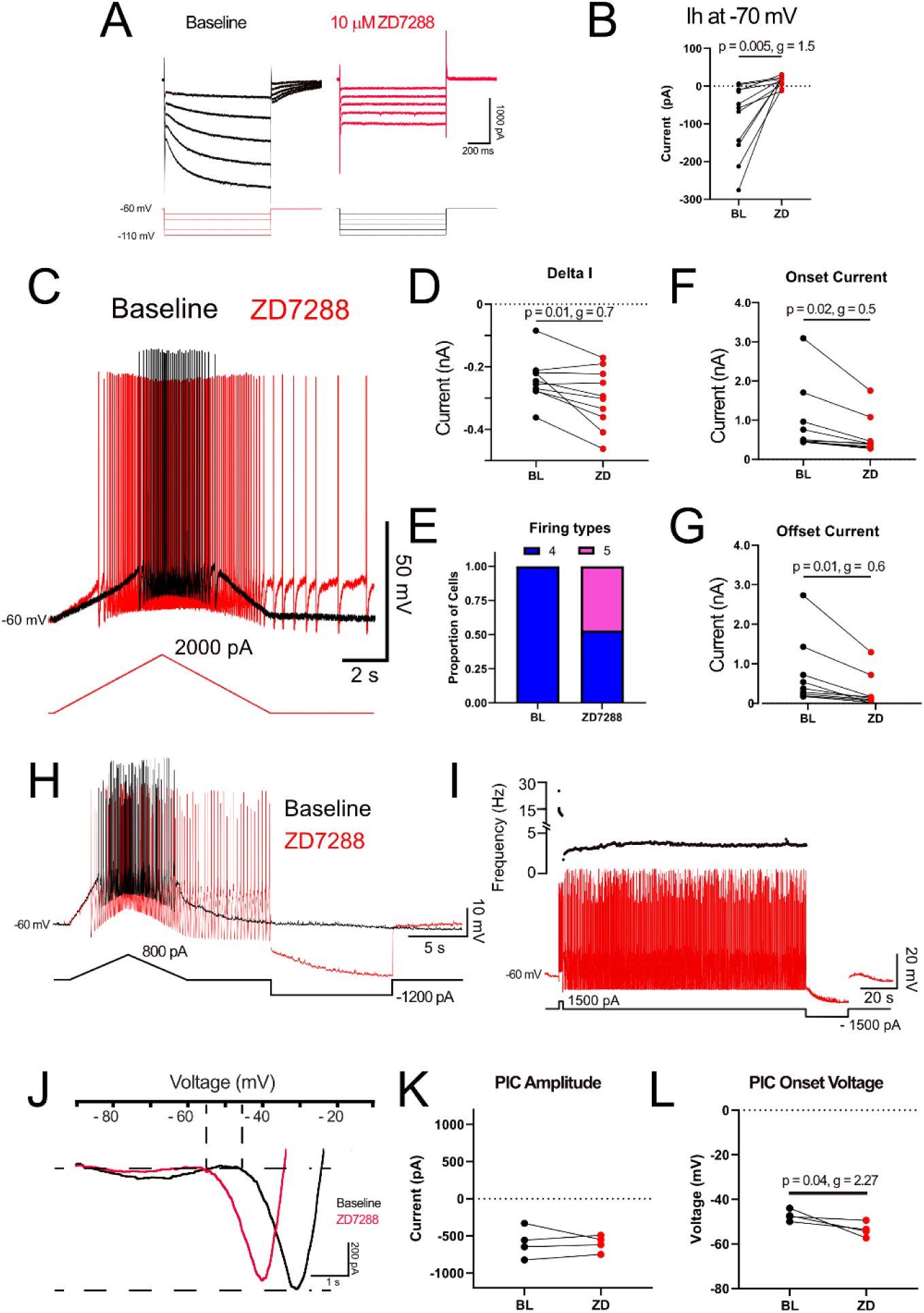
HCN channels prevent self sustained firing at the third postnatal week. (A) Representative voltage clamp traces illustrating the H-current measured in response to a 10-mV hyperpolarizing voltage step applied from -60 mV before (black) and after application of the HCN channel blocker ZD7288 (10 µM) in 10 delayed firing motoneurons obtained from mice during the third postnatal week (P14-20). (B) ZD7288 eliminates H current measured at -70 mV. (C) Representative traces of the membrane potential and repetitive firing during a triangular depolarizing current ramp before (black) and after (red) application of ZD7288. ZD7288 increased recruitment-derecruitment hysteresis (D; Delta I) and produced a shift from Type 4 firing hysteresis in all cells to the emergence of a 5^th^ firing type characterized by self sustained firing (E). ZD7288 decreased the current at firing onset on the ascending limb of the ramp (F) decreased the current of firing offset on the descending limb of the ramp (G). Hyperpolarizing current was needed to terminate self-sustained firing elicited after triangular current ramps (H) or brief depolarizing current steps (I) in the presence of ZD7288. J) Representative traces of a leak-subtracted and filtered PIC measured in voltage clamp before (black) and after (red) application of the HCN channel blocker ZD7288 (10 uM; n = 4). ZD7288 did not change the amplitude (K) of the PIC but hyperpolarized the PIC onset voltage (L). Data are presented as individual data points and were analyzed using a paired t-test. P values are reported when significant differences were detected. Hedge’s g values indicated small (0.2-0.49), medium (0.5-0.79) and large (>0.8) effect sizes.

## Discussion

The present study examined the developmental maturation of persistent inward currents (PICs) and their relationship to non-linear firing dynamics leading to recruitment–derecruitment and firing hysteresis in lumbar spinal motoneurons of the postnatal mouse. Although PICs are widely assumed to be the primary determinant of sustained firing hysteresis in motoneurons (Hounsgaard & Kiehn, 1989; Li & Bennett, 2003; Heckman *et al*., 2008b), alternative mechanisms have not been tested directly. This issue is particularly important because recruitment–derecruitment hysteresis measured in human motor units is commonly quantified using ΔF from paired motor unit recordings during triangular isometric contractions and interpreted as an index of PIC amplitude or neuromodulatory drive (Gorassini *et al*., 2002; Heckmann *et al*., 2005; Heckman *et al*., 2008a, 2008b; Goodlich *et al*., 2024; Mesquita *et al*., 2024). Here we addressed three related hypotheses. First, that developmental increases in PIC amplitude underlie the emergence of recruitment–derecruitment hysteresis during the onset of hindlimb weight bearing (Clarac *et al*., 2004; Quinlan *et al*., 2011; Sharples & Miles, 2021). Second, that altering PIC amplitude through pharmacological manipulation of NaV1.6 and L-type calcium channels would produce predictable changes in firing hysteresis (Carlin *et al*., 2000; Li & Bennett, 2003; Li *et al*., 2004). Third, that other conductances—particularly those mediated by potassium and HCN channels—may shape hysteresis by modifying the balance between recruitment and derecruitment currents (Manuel *et al*., 2007; Bos *et al*., 2018; Sharples & Miles, 2021; Sharples *et al*., 2023; Harris-Warrick *et al*., 2024). The conceptual advance of this study is that it directly tests the long-standing assumption that PIC amplitude determines hysteretic firing behaviour.

Our results partially supported the first hypothesis but challenged the latter two. PIC amplitude and recruitment–derecruitment hysteresis increased in parallel in fast-type delayed firing motoneurons during the onset of weight bearing, whereas slow-type immediate firing motoneurons showed little developmental change. However, although NaV1.6 and L-type calcium channels contributed substantially to PIC amplitude, manipulating these currents did not alter recruitment–derecruitment or firing hysteresis (Bouhadfane *et al*., 2013; Sharples & Miles, 2021; Drouillas *et al*., 2023). Moreover, pharmacological manipulations that increased PIC amplitude did not produce the predicted increase in sustained firing hysteresis; notably, muscarine increased PIC amplitude while paradoxically reducing recruitment–derecruitment hysteresis and promoting adaptive firing hysteresis (Revill *et al*., 2019; Sharples *et al*., 2023). Instead, potassium channels—including KCNQ and Kv1.2—strongly shaped hysteretic firing by altering the balance of onset and offset currents during recruitment and derecruitment (Bos *et al*., 2018; Verneuil *et al*., 2020; Singh *et al*., 2025). Finally, HCN channels limited the emergence of bistable firing in fast motoneurons by the third postnatal week (Manuel *et al*., 2007; Sharples & Miles, 2021). Together these findings indicate that although PICs and firing hysteresis mature together during development, sustained hysteretic firing cannot be attributed solely to PIC amplitude. Instead, hysteresis emerges from interactions between inward and outward currents that shape recruitment–derecruitment asymmetry.

### Developmental emergence of PICs and hysteresis

Hindlimb weight-bearing is a pivotal behavioral milestone in rodents, emerging toward the end of the second postnatal week (Altman & Sudarshan, 1975; Brocard *et al*., 1999). It requires sustained activation of postural motor units to support antigravity function, and it has long been proposed that maturation of dendritic PICs underlies this transition by enabling motoneurons to produce sustained action potential discharge following transient excitatory inputs. Consistent with this view, we observed a developmental increase in both PIC amplitude and recruitment–derecruitment hysteresis specifically in delayed firing motoneurons during the onset of weight bearing. Despite the increase in PIC amplitude and delta I, the proportion of Types 1–4 firing hysteresis remained relatively stable across postnatal development. These findings agree with prior work showing that PICs become more prominent in motoneurons around the second postnatal week and contribute to enhanced excitability, however the changes in delta I before the end of the second week are relatively subtle (Quinlan *et al*., 2011; Sharples & Miles, 2021). However, a surprising feature of our results, which is in line with recent findings from Harris-Warrick and colleagues (Harris-Warrick *et al*., 2024), is that the developmental increase in PIC amplitude was most pronounced in delayed firing motoneurons, considered fast-type, rather than in immediate firing slow-type motoneurons that innervate fatigue-resistant units thought to be essential for postural tone. Human studies similarly report larger PIC estimates (ΔF) in higher threshold motor units and in muscles enriched with fast motor units, such as tibialis anterior, compared with muscles more involved in postural control such as soleus (Orssatto *et al*., 2021; Jenz *et al*., 2023; Škarabot *et al*., 2025). Although differences in firing rates between fast and slow muscle types could influence ΔF estimates in humans, our results suggest that the functional role of PICs extends beyond slow motor unit stabilization and may contribute to dynamic amplification or rapid recruitment strategies in fast units. Thus, while PICs increase during the period when weight-bearing emerges, their distribution across motoneuron subtypes challenges classical assumptions and calls for a reassessment of their role in postural versus phasic motor control.

### PIC amplitude does not dictate firing hysteresis

Although PIC amplitude and sustained firing hysteresis increased in parallel during development, our pharmacological experiments demonstrate that PIC amplitude alone does not determine sustained hysteretic firing. Blocking NaV1.6 or L-type calcium channels significantly reduced PIC amplitude but did not reduce impact firing hysteresis (Sharples & Miles, 2021; Drouillas *et al*., 2023). Similarly, co-blockade that reduced PIC amplitude by nearly 60% failed to change firing hysteresis in a predictable way. Conversely, muscarine nearly doubled PIC amplitude yet promoted adaptive firing hysteresis by selectively increasing offset current. Together these results indicate that hysteresis is not linearly—or even directly—related to PIC magnitude. This finding contrasts with the canonical model in which suggests that PIC activation drives sustained counterclockwise hysteresis in motoneuron firing (Hounsgaard *et al*., 1984; Hounsgaard & Kiehn, 1989; Hultborn *et al*., 2003; Heckman *et al*., 2003). This model is supported by classic studies in cats and turtles showing bistable firing and plateau potentials dependent on dendritic L-type calcium channels under strong neuromodulatory drive (Hounsgaard & Kiehn, 1989; Carlin *et al*., 2000; Hultborn *et al*., 2003; Bui *et al*., 2006; Elbasiouny *et al*., 2006; Heckman *et al*., 2008a). Our data suggest that in neonatal mouse motoneurons, particularly in slice preparations with reduced neuromodulatory tone, PIC amplitude alone is insufficient to explain recruitment–derecruitment nonlinearities. Instead, other ionic mechanisms appear to provide critical shaping of firing hysteresis (Manuel *et al*., 2007; Bos *et al*., 2018), which could account for examples where motoneurons produce positive delta I values and adaptive firing hysteresis despite the presence of robust PICs (Revill *et al*., 2019). One interpretation of these findings is that PICs provide the inward drive necessary for sustained firing but do not themselves determine the asymmetry between recruitment and derecruitment that defines hysteresis. Alternatively, computational studies inspired by experimental work in the stomatogastric ganglion have elegantly demonstrated that degenerate circuit mechanisms can produce similar network outputs, such that distinct combinations of intrinsic and synaptic conductances generate comparable firing patterns (Prinz *et al*., 2004; Mellen, 2008). These findings raise the possibility that similar degeneracy may exist in spinal motoneurons, where multiple ionic mechanisms could give rise to firing behaviors—such as hysteresis or sustained discharge—that are often interpreted as signatures of persistent inward currents.

### Potassium channels shape recruitment–derecruitment asymmetry

If PICs alone do not determine firing hysteresis, then other conductances that influence membrane potential near spike threshold may play a key role in shaping these firing behaviors. Potassium channels are strong candidates in this regard because they regulate excitability in the subthreshold voltage range (Deardorff *et al*., 2021) and are well positioned to differentially influence onset and offset currents during recruitment and derecruitment.

In the present study, we identified KCNQ and Kv1.2 channels as modulators of hysteretic firing. KCNQ channels, which conduct the M-type potassium current, are ideally positioned to counteract PICs due to their subthreshold activation and non-inactivating outward current (Brown & Adams, 1980; Verneuil *et al*., 2020; Singh *et al*., 2025; Gaudreau & Bui, 2026). We found that blocking KCNQ channels increased PIC amplitude but paradoxically reduced hysteresis, whereas activation of KCNQ channels enhanced hysteresis and shifted firing toward sustained counterclockwise patterns. These results highlight that outward currents, far from simply opposing inward currents, can create the conditions for recruitment–derecruitment hysteresis by differentially influencing onset versus offset currents for firing. Kv1.2 channels provide an additional complementary mechanism. Their rapid activation and slow inactivation underlie the delayed firing phenotype of fast motoneurons (Bos *et al*., 2018) and we hypothesized that their slow inactivation would support sustained firing on the descending limb of the ramp. While blocking Kv1.2 channels reduced delta I and shifted firing from sustained to more adaptive patterns as expected, it did so by decreasing onset current without impacting the offset current. Nevertheless, these results point to potassium currents at the axon initial segment as critical determinants of hysteretic firing. These findings align with growing evidence that motoneuron input–output nonlinearities are strongly influenced by potassium channel kinetics (Manuel *et al*., 2014; Leroy *et al*., 2015; Deutsch & Elbasiouny, 2024; Molkov *et al*., 2025). Importantly, these results shift the explanatory framework: rather than attributing firing hysteresis solely to dendritic PICs, our findings suggest that potassium channels—particularly those localized to the axon initial segment—shape recruitment–derecruitment asymmetry and therefore strongly influence the expression of sustained firing behaviors.

### HCN channels limit bistable firing

By the third postnatal week, fast motoneurons develop a resting H-current that delays their recruitment (Sharples & Miles, 2021). Strikingly, blocking HCN channels in the present study produced bistable firing in nearly half of fast motoneurons, an observation that is consistent with previous reports of increased bistability in resonant motoneurons - a key property medicated by HCN channels (Manuel *et al*., 2007). In a subset of cells, HCN channel blockade also hyperpolarized PIC onset voltage while producing minimal change in PIC amplitude, suggesting that HCN channels may act as a shunt conductance near resting membrane potential that delays PIC activation. It is therefore possible that reactivation of HCN channels during the descending limb of triangular current ramps prevents bistable firing and maintains the temporal fidelity of synaptic inputs to fast motoneurons. Together these observations suggest that PICs, potassium channels, and HCN channels act in concert to shape nonlinear motoneuron firing, with their relative influence depending on developmental stage, neuromodulatory state, and motoneuron subtype.

### Neuromodulation as a dynamic regulator of PIC and firing hysteresis

Neuromodulators play a critical role in shaping motoneuron excitability, influencing both normal motor function and the emergence of aberrant firing in disease states such as spasticity, amyotrophic lateral sclerosis, and cerebral palsy. Traditionally, their effects have been attributed to modulation of PICs. However, the actions of neuromodulators extend beyond PICs, affecting multiple ionic conductances that together are likely to define recruitment–derecruitment dynamics and firing hysteresis. Our muscarinic experiments illustrate this complexity. Muscarine robustly increased PIC amplitude yet paradoxically promoted adaptive firing hysteresis, an effect that can be explained, at least in part, by inhibition of KCNQ channels (Revill *et al*., 2019; Sharples *et al*., 2023). Consistent with this idea, pharmacological blockade of KCNQ channels similarly increased PIC amplitude and decreased delta I. Importantly, the underlying mechanisms differed: KCNQ channel blockade reduced delta I by selectively decreasing onset current without affecting offset current, whereas muscarine reduced delta I by increasing offset current without altering onset. This divergence highlights that neuromodulators rarely act on a single channel type; instead, they simultaneously modify multiple conductances, producing net effects that may diverge from simple predictions based on PIC amplitude alone (Marder, 2011; Perrier *et al*., 2013; Marder *et al*., 2014; Sharples *et al*., 2014; Nascimento *et al*., 2020). Additional targets of neuromodulatory regulation likely include the sodium–potassium ATPase, which exerts activity-dependent inhibition of motoneuron excitability (Picton *et al*., 2017; Hachoumi *et al*., 2022; Akkuratov *et al*., 2025; Sharples *et al*., 2025), which can be modulated by acetylcholine in the brain (Tiwari *et al*., 2018; Mohan *et al*., 2019, 2021), and is a key target of neuromodulators in spinal networks (Picton *et al*., 2017; Hachoumi *et al*., 2022). It is therefore a reasonable hypothesis that increasing sodium pump activity would promote adaptive firing hysteresis. In vivo, serotonergic, cholinergic, and noradrenergic systems converge to regulate both inward and outward currents, enabling flexible control of motoneuron gain and persistent firing in a behaviorally relevant context (Heckman *et al*., 2008a; Goodlich *et al*., 2023, 2024). Our results are particularly important in the context of injury and disease where PICs and their neuromodulatory control have been implicated in the generation of spasticity and motor dysfunction (Murray *et al*., 2010; ElBasiouny *et al*., 2010; Quinlan *et al*., 2011; D’Amico *et al*., 2013; Brocard *et al*., 2016; Steele *et al*., 2020; Marcantoni *et al*., 2020; Jiang *et al*., 2021; Reedich *et al*., 2023; Delestrée *et al*., 2023; Deutsch & Elbasiouny, 2024; Pagiazitis *et al*., 2025). These observations therefore set the stage for our findings and future studies, which identify novel ionic mechanisms—including potassium and HCN channels—that contribute to neuromodulatory control and may underlie dysfunction in injury and disease.

### Methodological considerations

Several caveats should be considered when interpreting our results. Slice preparations limit dendritic integrity and neuromodulatory input, potentially underestimating the contribution of dendritic PICs (Mousa & Elbasiouny, 2021). Somatic current injection may also insufficiently activate distal channels that are robustly engaged by synaptic inputs in vivo. Recordings were also conducted at sub physiological temperature, which can not only alter channel kinetics (Bouhadfane *et al*., 2013), an important consideration given that temperature-sensitive TRPM5 channels are critical for generating bistability (Bos *et al*., 2021). Starting membrane potential also influences the likelihood of self-sustained discharge (Bouhadfane *et al*., 2013; Mahrous *et al*., 2024; Molkov *et al*., 2025). Nevertheless, we still observed bistable firing at these temperatures following HCN channel blockade from a starting membrane potential of -60 mV. Together, these factors indicate that the relative contributions of PICs and outward conductances may vary under more physiological conditions but do not negate our conclusions. Finally, developmental comparisons were made in transverse slices; complementary in vivo recordings could confirm whether the changes we observed translate into functional differences during natural motor behavior (de Lourdes Martínez-Silva *et al*., 2025).

### Functional implications

Our findings have important implications for understanding motor unit behavior in humans, where ΔF derived from paired motor unit recordings is widely interpreted as an estimate of PIC amplitude or neuromodulatory state (Powers *et al*., 2008; Udina *et al*., 2010; Revill & Fuglevand, 2011; Vandenberk & Kalmar, 2014; Powers & Heckman, 2015). Similarly, several other electrophysiological signatures commonly interpreted as indicators of PIC activation may also arise from the combined influence of multiple conductances. For example, analyses of motor unit firing-rate trajectories, including the “brace-height” metric used to quantify nonlinear acceleration of discharge during voluntary contractions, have been interpreted as evidence of PIC recruitment in human motoneurons (Beauchamp *et al*., 2023; Škarabot *et al*., 2025). Further, quantification of secondary and tertiary firing ranges in motoneuron and motor unit firing has also been used to infer the engagement of intrinsic depolarizing conductances that accelerate discharge once firing is established (Lee & Heckman, 1998; Bennett *et al*., 1998; Meehan *et al*., 2010a; Binder *et al*., 2020; Afsharipour *et al*., 2020). Finally, intracellular recordings in animal models frequently describe a subthreshold acceleration of membrane potential preceding spike threshold, which has likewise been attributed to the activation of inward currents (Kuo *et al*., 2006; Delestrée *et al*., 2014; Jensen *et al*., 2020; Sharples & Miles, 2021). However, the specific ionic mechanisms underlying these features have not been directly tested, and it remains unclear to what extent PICs versus other voltage-dependent conductances contribute to these dynamics. Together, these observations suggest that many physiological proxies used to estimate PIC influence on motoneuron output—including ΔF, nonlinear firing-rate acceleration, and secondary-range discharge behavior—may reflect the emergent behavior of interacting channel populations rather than the action of a single dominant inward current. Our results suggest that ΔF, and other non-linear firing properties, may reflect the integrated contribution of multiple intrinsic and neuromodulatory mechanisms—including potassium and HCN channel dynamics—that shape recruitment–derecruitment asymmetry.

## Conclusion

In summary, this study demonstrates that while PIC amplitude and recruitment–derecruitment hysteresis increase together during postnatal development, PIC magnitude alone does not determine sustained hysteretic firing. Instead, PICs may provide the inward drive required for sustained firing, but potassium and HCN channels determine whether hysteretic firing actually emerges. These findings expand the traditional PIC-centric view of motoneuron bistability and suggest that sustained firing behaviors arise from the interaction of multiple inward and outward conductances. This broader framework also has important implications for the interpretation of ΔF measurements in human motor unit studies, where hysteresis is often treated as a proxy for PIC amplitude or neuromodulatory state. From our work, we suggest that ΔF may reflect the integrated contribution of several intrinsic mechanisms that shape motoneuron input–output nonlinearities.

## Additional Information

### Contributions

Study Conception and Design: SAS, GBM; Data acquisition and analysis: SAS; Preparation of Figures: SAS; First draft and revision of manuscript: SAS, GBM; All authors approved the final version of the manuscript.

## Acknowledgements

The authors reserve the right to apply a Creative Commons Attribution (CC BY) licence to any Author Accepted manuscript version arising from this submission.

## Data Availability Statement

The research data supporting this publication will be made freely available in an open access data repository following acceptance to a peer-reviewed journal.

## Methods

### Animals

This study included unpublished data in addition to reanalysis of experimental data summarised in (Sharples & Miles, 2021), which included experiments performed on tissue obtained from 117 (male: n = 55; and female: n = 62) wild type C57Bl/6J mice at postnatal days (P) 7-20. We also present unpublished data obtained from 21 C57Bl/6J mice (P7-13; n = 9 male, n = 12 female) that were included in (Sharples *et al*., 2023). This study also included new experimental data obtained from 29 (male: n = 17; and female: n = 12) wild type C57Bl/6J mice at postnatal days P7-13. All procedures were conducted in accordance with the UK Animals (Scientific Procedures) Act 1986. Experiments conducted at the University of St Andrews were approved by the University of St Andrews Animal Welfare Ethics Committee and were covered under a Project Licence (PP8253850) approved by the Home Office. All animals were provided with unrestricted access to food and water and housed in climate-controlled conditions.

### Tissue preparation

Animals were sourced from an in-house colony within the St Mary’s Animal Unit at the University of St Andrews. Animals were killed using Schedule 1 procedures defined by the Home Office by performing a cervical dislocation followed by rapid decapitation. Animals were then eviscerated and pinned ventral side up in a dissecting chamber lined with silicone elastomer (Sylguard), filled with ice-cold (1-2 degrees Celsius) potassium gluconate based dissecting/slicing aCSF (containing in mM: 130 K-gluconate, 15 KCl, 0.05 EGTA, 20 HEPES, 25 D-glucose, 3 kynurenic acid, 2 Na-pyruvate, 3 myo-inositol, 1 Na-L-ascorbate; pH 7.4, adjusted with NaOH; osmolarity approximately 345 mOsm) that was continuously bubbled with carbogen (95% oxygen, 5% carbon dioxide). Spinal cords were exposed by performing a ventral vertebrectomy, cutting the dorsal roots and gently lifting the spinal cord from the spinal column. Spinal cords were removed within 3 - 5 minutes following cervical dislocation. Spinal cords were secured directly to an agar block (3 % agar) with VetBond surgical glue (3M) and glued to the base of the slicing chamber with cyanoacrylate adhesive. The tissue was immersed in ice-cold dissecting/slicing aCSF and bubbled with carbogen. Blocks of frozen slicing solution were also placed in the slicing chamber to keep the solution around 1-2 degrees Celsius. On average, the first slice was obtained within 10 minutes of decapitation which increased the likelihood of obtaining viable motoneurons in slices. 300 µm transverse slices were cut at a speed of 10 um/s on the vibratome (Leica VT1200) to minimize tissue compression during slicing. 3-4 slices were obtained from each animal. Slices were transferred to a recovery chamber filled with carbogenated pre-warmed (35 degrees Celsius) recovery aCSF (containing in mM: 119 NaCl, 1.9 KCl, 1.2 NaH2PO4, 10 MgSO4, 1 CaCl, 26 NaHCO3, 20 glucose, 1.5 kynurenic acid, 3% dextran) for thirty minutes after completion of the last slice which took 10 -15 minutes on average. Following recovery, slices were transferred to a chamber filled with warm (35 degrees Celsius) recording aCSF (containing in mM: 127 NaCl, 3 KCl, 1.25 NaH2PO4, 1 MgCl, 2 CaCl2, 26 NaHCO3, 10 glucose), bubbled with carbogen, and allowed to equilibrate at room temperature (maintained at 23-25 degrees Celsius) for at least one hour before experiments were initiated.

### Whole cell patch clamp electrophysiology

This study includes data from whole-cell patch clamp recordings obtained from a total of 291 lumbar motoneurons. 219 of these cells were reanalyzed using data published in (Sharples & Miles, 2021). We also present novel data from motoneurons that were studied in (Sharples *et al*., 2023) and included new experiments on 41 additional motoneurons. In these experiments, spinal cord slices were stabilized in a recording chamber with fine fibres secured to a platinum harp and visualized with a 40x objective with infrared illumination and differential interference contrast (DIC) microscopy. A large proportion of the motoneurons studied were identified based on location in the ventrolateral region with somata greater than 20 µm. Recordings were obtained from a subset of motoneurons that had been retrogradely labelled with Fluorogold (Fluorochrome, Denver, CO). Fluorogold was dissolved in sterile saline solution and 0.04 mg/g injected intraperitoneally 24-48 hours prior to experiments(Miles *et al*., 2005). In addition to recording from larger FG-positive cells, this approach allowed us to more confidently target smaller motoneurons. Motoneurons were visualized and whole cell recordings obtained under DIC illumination with pipettes (L: 100 mm, OD: 1.5 mm, ID: 0.84 mm; World Precision Instruments) pulled on a Flaming Brown micropipette puller (Sutter instruments P97) to a resistance of 2.5-3.5 MΩ. Pipettes were back-filled with intracellular solution (containing in mM: 140 KMeSO4, 10 NaCl, 1 CaCl2, 10 HEPES, 1 EGTA, 3 Mg-ATP and 0.4 GTP-Na2; pH 7.2-7.3, adjusted with KOH).

Signals were amplified and filtered (6 kHz low pass Bessel filter) with a Multiclamp 700 B amplifier, acquired at 20 kHz using a Digidata 1440A digitizer with pClamp Version 10.7 software (Molecular Devices) and stored on a computer for offline analysis.

### Identification of fast and slow motoneuron types

Motoneuron subtypes were identified using a protocol established by (Leroy *et al*., 2014), which differentiates motoneuron type based on the latency to the first spike when injecting a 5 second square depolarizing current near the threshold for repetitive firing. Using this approach we were able to identify 2 main firing profiles - a delayed repetitive firing profile with accelerating spike frequency, characteristic of fast-type motoneurons, and an immediate firing profile with little change in spike frequency, characteristic of slow-type motoneurons (Figure 1).

All motoneuron intrinsic properties were studied by applying a bias current to maintain the membrane potential at -60 mV. Values reported are not liquid junction potential corrected to facilitate comparisons with previously published data (Miles *et al*., 2007; Quinlan *et al*., 2011; Durand *et al*., 2015; Nascimento *et al*., 2020, 2024; Smith & Brownstone, 2020; Özyurt *et al*., 2022; Pocratsky *et al*., 2023). Cells were excluded from analysis if access resistance was greater than 20 MΩ or changed by more than 5 MΩ over the duration of the recording, or if spike amplitude measured from threshold (described below) was less than 60 mV.

### Pharmacology

Nifedipine (20 µM; Tocris) was used to assess the contribution of L-type calcium channels, and 4,9 - Anhydro Tetrodotoxin (200 nM; Tocris) to assess the contribution of NaV1.6 channels to PIC and hysteresis. KCNQ channels that underlie the M current were activated with ICA069673 (ICA73: 10 uM; Tocris) or retigabine (10 uM; Tocris) or blocked with XE991 (10 uM; Tocris). Kv1.2 channels were blocked with Tityustoxin (800 nM; Alomone). HCN Channels that underlie the H-current were blocked with ZD7288 (10 uM; Tocris).

### Data acquisition and analysis

Passive properties including capacitance, membrane time constant (tau), and input resistance (Ri) were measured during a hyperpolarizing current pulse that brought the membrane potential from -60 to -70mV. Input resistance was measured from the initial voltage trough to minimize the impact of slower acting active conductances (eg. Ih, sag). The time constant was measured as the time it took to reach 2/3 of the peak voltage change. Capacitance was calculated by dividing the time constant by the input resistance (C=T/R).

PICs were measured in voltage clamp during slow depolarizing voltage ramps (10 mV/s from -90 to -10 mV) over 8 seconds (Quinlan *et al*., 2011; Steele *et al*., 2020; Huh *et al*., 2021). PIC onset voltage and peak current amplitude measured from post-hoc leak-subtracted traces as in previous studies (Quinlan *et al*., 2011; Steele *et al*., 2020; Verneuil *et al*., 2020).

Recruitment-derecruitment and firing hysteresis were measured in current clamp using triangular, depolarizing current ramps with 5 second rise and fall times (Bennett *et al*., 2001; Li & Bennett, 2003; Durand *et al*., 2015; Steele *et al*., 2020). Triangular depolarizing current ramps were set to a peak current of 2 times repetitive firing threshold current (determined with a 100pA/s depolarizing current ramp initiated from -60 mV). Recruitment-derecruitment hysteresis was measured by calculating the difference (delta I) between the current at firing onset on the ascending component of the ramp and the current at derecruitment on the descending component of the ramp. Firing hysteresis was also assessed by examining the frequency-current trajectories on the ascending and descending components of the ramp. We subdivided cells into 1 of 4 types based on previously-defined criteria identifying the pattern of firing hysteresis on ascending and descending portions of the ramp (Bennett *et al*., 2001) (Type 1: Linear, Type 2: Adapting clockwise hysteresis, Type 3: Linear Sustained, Type 4: Accelerating sustained Counter-clockwise hysteresis). In addition, we also identified a fifth firing type following blockade of HCN channels, characterized by sustained firing that continued beyond the end of the descending portion of the triangular current ramp. In this fifth firing type, hyperpolarizing current was required to terminate sustained firing.

### Research design and statistical analysis

Two factor analysis of variance (ANOVA) was performed to study changes in PIC and recruitment-derecruitment hysteresis (delta I) in motoneuron subtypes across developmental time points or to determine effects of pharmacological agents at different developmental stages. Paired or unpaired t-tests were performed when comparing two conditions. Firing hysteresis (Types 1-4) were analyzed with either a Kruskal-Wallis test when comparing more than two conditions or Wilcoxon test when comparing two conditions. Appropriate and equivalent nonparametric tests (Mann-Whitney or Kruskal-Wallis) were conducted when data failed tests of normality or equal variance with Shapiro Wilk and Brown-Forsythe tests, respectively. Statistical tests were complemented with Hedge’s g to indicate effect sizes, categorized as small (0.2-0.49), medium (0.5-0.79) and large (>0.8). Individual data points for all cells are presented in figures with mean ± SD. Statistical analyses were performed using Graph Pad Version 9.0 (Prism, San Diego, CA, USA). All statistical tests and results are summarized in Table 1 and are annotated in text where appropriate.

## References

Afsharipour B, Manzur N, Duchcherer J, Fenrich KF, Thompson CK, Negro F, Quinlan KA, Bennett DJ & Gorassini MA (2020). Estimation of self-sustained activity produced by persistent inward currents using firing rate profiles of multiple motor units in humans. J Neurophysiol 124, 63–85.

Akkuratov EE, Sorrell F, Picton LD, Sousa VC, Paucar M, Jans D, Svensson L-B, Lindskog M, Fritz N, Liebmann T, Sillar KT, Rosewich H, Svenningsson P, Brismar H, Miles GB & Aperia A (2025). ATP1A3 dysfunction causes motor hyperexcitability and afterhyperpolarization loss in a dystonia model. Brain 148, 1099–1105.

Alaburda A, Perrier J-F & Hounsgaard J (2002). An M-like outward current regulates the excitability of spinal motoneurones in the adult turtle. J Physiol 540, 875–881.

Altman J & Sudarshan K (1975). Postnatal development of locomotion in the laboratory rat. Anim Behav 23, 896–920.

Beauchamp JA, Pearcey GEP, Khurram OU, Chardon M, Wang YC, Powers RK, Dewald JPA & Heckman CJ (2023). A geometric approach to quantifying the neuromodulatory effects of persistent inward currents on individual motor unit discharge patterns. J Neural Eng; DOI: 10.1088/1741-2552/acb1d7.

Bennett DJ, Hultborn H, Fedirchuk B & Gorassini M (1998). Synaptic activation of plateaus in hindlimb motoneurons of decerebrate cats. J Neurophysiol 80, 2023–2037.

Bennett DJ, Li Y & Siu M (2001). Plateau potentials in sacrocaudal motoneurons of chronic spinal rats, recorded in vitro. J Neurophysiol 86, 1955–1971.

Binder MD, Powers RK & Heckman CJ (2020). Nonlinear Input-Output Functions of Motoneurons. Physiology (Bethesda) 35, 31–39.

Bos R, Drouillas B, Bouhadfane M, Pecchi E, Trouplin V, Korogod SM & Brocard F (2021). Trpm5 channels encode bistability of spinal motoneurons and ensure motor control of hindlimbs in mice. Nat Commun 12, 6815.

Bos R, Harris-Warrick RM, Brocard C, Demianenko LE, Manuel M, Zytnicki D, Korogod SM & Brocard F (2018). Kv1.2 Channels Promote Nonlinear Spiking Motoneurons for Powering Up Locomotion. Cell Rep 22, 3315–3327.

Bouhadfane M, Tazerart S, Moqrich A, Vinay L & Brocard F (2013). Sodium-mediated plateau potentials in lumbar motoneurons of neonatal rats. J Neurosci 33, 15626–15641.

Brocard C, Plantier V, Boulenguez P, Liabeuf S, Bouhadfane M, Viallat-Lieutaud A, Vinay L & Brocard F (2016). Cleavage of Na(+) channels by calpain increases persistent Na(+) current and promotes spasticity after spinal cord injury. Nat Med 22, 404–411.

Brocard F, Vinay L & Clarac F (1999). Development of hindlimb postural control during the first postnatal week in the rat. Brain Res Dev Brain Res 117, 81–89.

Brown DA & Adams PR (1980). Muscarinic suppression of a novel voltage-sensitive K+ current in a vertebrate neurone. Nature 283, 673–676.

Bui TV, Ter-Mikaelian M, Bedrossian D & Rose PK (2006). Computational estimation of the distribution of L-type Ca(2+) channels in motoneurons based on variable threshold of activation of persistent inward currents. J Neurophysiol 95, 225–241.

Button D (2008). The Biophysical Properties of Rat Hindlimb Motoneurones Before and After the Removal of Descending, Ascending, and Afferent Inputs.

Button DC, Kalmar JM, Gardiner K, Cahill F & Gardiner PF (2007). Spike frequency adaptation of rat hindlimb motoneurons. J Appl Physiol (1985) 102, 1041–1050.

Carlin KP, Jones KE, Jiang Z, Jordan LM & Brownstone RM (2000). Dendritic L-type calcium currents in mouse spinal motoneurons: implications for bistability. Eur J Neurosci 12, 1635–1646.

Clarac F, Brocard F & Vinay L (2004). The maturation of locomotor networks. Prog Brain Res 143, 57–66.

Conway BA, Hultborn H, Kiehn O & Mintz I (1988). Plateau potentials in alpha-motoneurones induced by intravenous injection of L-dopa and clonidine in the spinal cat. J Physiol 405, 369–384.

D’Amico JM, Murray KC, Li Y, Chan KM, Finlay MG, Bennett DJ & Gorassini MA (2013). Constitutively active 5-HT2/α1 receptors facilitate muscle spasms after human spinal cord injury. J Neurophysiol 109, 1473–1484.

Deardorff AS, Romer SH & Fyffe REW (2021). Location, location, location: the organization and roles of potassium channels in mammalian motoneurons. J Physiol 599, 1391–1420.

Delestrée N, Manuel M, Iglesias C, Elbasiouny SM, Heckman CJ & Zytnicki D (2014). Adult spinal motoneurones are not hyperexcitable in a mouse model of inherited amyotrophic lateral sclerosis. J Physiol 592, 1687–1703.

Delestrée N, Semizoglou E, Pagiazitis JG, Vukojicic A, Drobac E, Paushkin V & Mentis GZ (2023). Serotonergic dysfunction impairs locomotor coordination in spinal muscular atrophy. Brain 146, 4574–4593.

Deutsch AJ & Elbasiouny SM (2024). Dysregulation of persistent inward and outward currents in spinal motoneurons of symptomatic SOD1-G93A mice. J Physiol 602, 3715–3736.

Drouillas B, Brocard C, Zanella S, Bos R & Brocard F (2023). Persistent Nav1.1 and Nav1.6 currents drive spinal locomotor functions through nonlinear dynamics. Cell Rep 42, 113085.

Durand J, Filipchuk A, Pambo-Pambo A, Amendola J, Borisovna Kulagina I & Guéritaud J-P (2015). Developing electrical properties of postnatal mouse lumbar motoneurons. Front Cell Neurosci 9, 349.

Elbasiouny SM, Bennett DJ & Mushahwar VK (2006). Simulation of Ca2+ persistent inward currents in spinal motoneurones: mode of activation and integration of synaptic inputs. J Physiol 570, 355–374.

ElBasiouny SM, Schuster JE & Heckman CJ (2010). Persistent inward currents in spinal motoneurons: important for normal function but potentially harmful after spinal cord injury and in amyotrophic lateral sclerosis. Clin Neurophysiol 121, 1669–1679.

Gaudreau SF & Bui TV (2026). Distinct developmental dynamics of opposing persistent currents shape motoneuron firing during motor maturation of zebrafish. J Physiol 604, 2083–2109.

Ghezzi F, Corsini S & Nistri A (2017). Electrophysiological characterization of the M-current in rat hypoglossal motoneurons. Neuroscience 340, 62–75.

Ghezzi F, Monni L & Nistri A (2018). Functional up-regulation of the M-current by retigabine contrasts hyperexcitability and excitotoxicity on rat hypoglossal motoneurons. J Physiol 596, 2611–2629.

Goodlich BI, Del Vecchio A, Horan SA & Kavanagh JJ (2023). Blockade of 5-HT receptors suppresses motor unit firing and estimates of persistent inward currents during voluntary muscle contraction in humans. J Physiol 601, 1121–1138.

Goodlich BI, Pearcey GEP, Del Vecchio A, Horan SA & Kavanagh JJ (2024). Antagonism of 5-HT receptors attenuates self-sustained firing of human motor units. J Physiol 602, 1759–1774.

Gorassini M, Yang JF, Siu M & Bennett DJ (2002). Intrinsic activation of human motoneurons: reduction of motor unit recruitment thresholds by repeated contractions. J Neurophysiol 87, 1859–1866.

Hachoumi L, Rensner R, Richmond C, Picton L, Zhang H & Sillar KT (2022). Bimodal modulation of short-term motor memory via dynamic sodium pumps in a vertebrate spinal cord. Curr Biol 32, 1038–1048.e2.

Hamm TM, Turkin VV, Bandekar NK, O’Neill D & Jung R (2010). Persistent currents and discharge patterns in rat hindlimb motoneurons. J Neurophysiol 104, 1566–1577.

Harris-Warrick RM, Pecchi E, Drouillas B, Brocard F & Bos R (2023). A size principle for bistability in mouse spinal motoneurons. bioRxiv; DOI: 10.1101/2023.09.29.559784. Available at: 10.1101/2023.09.29.559784.

Harris-Warrick RM, Pecchi E, Drouillas B, Brocard F & Bos R (2024). Effect of size on expression of bistability in mouse spinal motoneurons. J Neurophysiol 131, 577–588.

Harvey PJ, Li X, Li Y & Bennett DJ (2006a). 5-HT2 receptor activation facilitates a persistent sodium current and repetitive firing in spinal motoneurons of rats with and without chronic spinal cord injury. J Neurophysiol 96, 1158–1170.

Harvey PJ, Li Y, Li X & Bennett DJ (2006b). Persistent sodium currents and repetitive firing in motoneurons of the sacrocaudal spinal cord of adult rats. J Neurophysiol 96, 1141–1157.

Heckman CJ, Hyngstrom AS & Johnson MD (2008a). Active properties of motoneurone dendrites: diffuse descending neuromodulation, focused local inhibition. J Physiol 586, 1225–1231.

Heckman CJ, Johnson M, Mottram C & Schuster J (2008b). Persistent inward currents in spinal motoneurons and their influence on human motoneuron firing patterns. Neuroscientist 14, 264–275.

Heckman CJ, Lee RH & Brownstone RM (2003). Hyperexcitable dendrites in motoneurons and their neuromodulatory control during motor behavior. Trends Neurosci 26, 688–695.

Heckmann CJ, Gorassini MA & Bennett DJ (2005). Persistent inward currents in motoneuron dendrites: implications for motor output. Muscle Nerve 31, 135–156.

Hounsgaard J, Hultborn H, Jespersen B & Kiehn O (1984). Intrinsic membrane properties causing a bistable behaviour of alpha-motoneurones. Exp Brain Res 55, 391–394.

Hounsgaard J, Hultborn H, Jespersen B & Kiehn O (1988). Bistability of alpha-motoneurones in the decerebrate cat and in the acute spinal cat after intravenous 5-hydroxytryptophan. J Physiol 405, 345–367.

Hounsgaard J & Kiehn O (1989). Serotonin-induced bistability of turtle motoneurones caused by a nifedipine-sensitive calcium plateau potential. J Physiol 414, 265–282.

Huh S, Heckman CJ & Manuel M (2021). Time Course of Alterations in Adult Spinal Motoneuron Properties in the SOD1(G93A) Mouse Model of ALS. eNeuro; DOI: 10.1523/ENEURO.0378-20.2021.

Huh S, Siripuram R, Lee RH, Turkin VV, O’Neill D, Hamm TM, Heckman CJ & Manuel M (2017). PICs in motoneurons do not scale with the size of the animal: a possible mechanism for faster speed of muscle contraction in smaller species. J Neurophysiol 118, 93–102.

Hultborn H, Denton ME, Wienecke J & Nielsen JB (2003). Variable amplification of synaptic input to cat spinal motoneurones by dendritic persistent inward current. J Physiol 552, 945–952.

Jean-Xavier C, Sharples SA, Mayr KA, Lognon AP & Whelan PJ (2018). Retracing your footsteps: developmental insights to spinal network plasticity following injury. J Neurophysiol 119, 521–536.

Jensen DB, Kadlecova M, Allodi I & Meehan CF (2020). Spinal motoneurones are intrinsically more responsive in the adult G93A SOD1 mouse model of amyotrophic lateral sclerosis. J Physiol 598, 4385–4403.

Jenz ST, Beauchamp JA, Gomes MM, Negro F, Heckman CJ & Pearcey GEP (2023). Estimates of persistent inward currents in lower limb motoneurons are larger in females than in males. J Neurophysiol 129, 1322–1333.

Jiang MC, Birch DV, Heckman CJ & Tysseling VM (2021). The Involvement of Ca1.3 Channels in Prolonged Root Reflexes and Its Potential as a Therapeutic Target in Spinal Cord Injury. Front Neural Circuits 15, 642111.

Kalmar JM, Button DC, Gardiner K, Cahill F & Gardiner PF (2009). Caloric restriction does not offset age-associated changes in the biophysical properties of motoneurons. J Neurophysiol 101, 548–557.

Kuo JJ, Lee RH, Zhang L & Heckman CJ (2006). Essential role of the persistent sodium current in spike initiation during slowly rising inputs in mouse spinal neurones. J Physiol 574, 819–834.

Lee RH & Heckman CJ (1998). Bistability in spinal motoneurons in vivo: systematic variations in persistent inward currents. J Neurophysiol 80, 583–593.

Leroy F, Lamotte d’Incamps B, Imhoff-Manuel RD & Zytnicki D (2014). Early intrinsic hyperexcitability does not contribute to motoneuron degeneration in amyotrophic lateral sclerosis. Elife; DOI: 10.7554/eLife.04046.

Leroy F, Lamotte d’Incamps B & Zytnicki D (2015). Potassium currents dynamically set the recruitment and firing properties of F-type motoneurons in neonatal mice. J Neurophysiol 114, 1963–1973.

Li X & Bennett DJ (2007). Apamin-sensitive calcium-activated potassium currents (SK) are activated by persistent calcium currents in rat motoneurons. J Neurophysiol 97, 3314–3330.

Li Y & Bennett DJ (2003). Persistent sodium and calcium currents cause plateau potentials in motoneurons of chronic spinal rats. J Neurophysiol 90, 857–869.

Li Y, Gorassini MA & Bennett DJ (2004). Role of persistent sodium and calcium currents in motoneuron firing and spasticity in chronic spinal rats. J Neurophysiol 91, 767–783.

de Lourdes Martínez-Silva M, Ahorklo RM, Reedich EJ, Imhoff-Manuel RD, Katenka N & Manuel M (2025). Machine-Learning Classification of Motor Unit Types in the Adult Mouse. bioRxiv; DOI: 10.1101/2025.11.18.689075. Available at: 10.1101/2025.11.18.689075.

MacDonell CW, Power KE, Chopek JW, Gardiner KR & Gardiner PF (2015). Extensor motoneurone properties are altered immediately before and during fictive locomotion in the adult decerebrate rat. J Physiol 593, 2327–2342.

Mahrous A, Birch D, Heckman CJ & Tysseling V (2024). Muscle Spasms after Spinal Cord Injury Stem from Changes in Motoneuron Excitability and Synaptic Inhibition, Not Synaptic Excitation. J Neurosci; DOI: 10.1523/JNEUROSCI.1695-23.2023.

Manuel M, Iglesias C, Donnet M, Leroy F, Heckman CJ & Zytnicki D (2009). Fast kinetics, high-frequency oscillations, and subprimary firing range in adult mouse spinal motoneurons. J Neurosci 29, 11246–11256.

Manuel M, Meunier C, Donnet M & Zytnicki D (2007). Resonant or not, two amplification modes of proprioceptive inputs by persistent inward currents in spinal motoneurons. J Neurosci 27, 12977–12988.

Manuel M, Zytnicki D & Meunier C (2014). The dendritic location of the L-type current and its deactivation by the somatic AHP current both contribute to firing bistability in motoneurons. Front Comput Neurosci 8, 4.

Marcantoni M, Fuchs A, Löw P, Bartsch D, Kiehn O & Bellardita C (2020). Early delivery and prolonged treatment with nimodipine prevents the development of spasticity after spinal cord injury in mice. Sci Transl Med; DOI: 10.1126/scitranslmed.aay0167.

Marder E (2011). Variability, compensation, and modulation in neurons and circuits. Proc Natl Acad Sci U S A 108 **Suppl 3**, 15542–15548.

Marder E, O’Leary T & Shruti S (2014). Neuromodulation of circuits with variable parameters: single neurons and small circuits reveal principles of state-dependent and robust neuromodulation. Annu Rev Neurosci 37, 329–346.

Meehan CF, Moldovan M, Marklund SL, Graffmo KS, Nielsen JB & Hultborn H (2010a). Intrinsic properties of lumbar motor neurones in the adult G127insTGGG superoxide dismutase-1 mutant mouse in vivo: evidence for increased persistent inward currents. Acta Physiol (Oxf) 200, 361–376.

Meehan CF, Sukiasyan N, Zhang M, Nielsen JB & Hultborn H (2010b). Intrinsic properties of mouse lumbar motoneurons revealed by intracellular recording in vivo. J Neurophysiol 103, 2599–2610.

Mellen NM (2008). Belt-and-suspenders as a biological design principle. Adv Exp Med Biol 605, 99–103.

Mesquita RNO, Taylor JL, Heckman CJ, Trajano GS & Blazevich AJ (2024). Persistent inward currents in human motoneurons: emerging evidence and future directions. J Neurophysiol 132, 1278–1301.

Miles GB, Dai Y & Brownstone RM (2005). Mechanisms underlying the early phase of spike frequency adaptation in mouse spinal motoneurones. J Physiol 566, 519–532.

Miles GB, Hartley R, Todd AJ & Brownstone RM (2007). Spinal cholinergic interneurons regulate the excitability of motoneurons during locomotion. Proc Natl Acad Sci U S A 104, 2448–2453.

Mohan S, Tiwari MN, Biala Y & Yaari Y (2019). Regulation of Neuronal Na/K-ATPase by Specific Protein Kinases and Protein Phosphatases. J Neurosci 39, 5440–5451.

Mohan S, Tiwari MN, Stanojević M, Biala Y & Yaari Y (2021). Muscarinic regulation of the neuronal Na /K -ATPase in rat hippocampus. J Physiol 599, 3735–3754.

Molkov YI, Krust F, Jeter R, Stell T, Mohammed MAY, Brocard F & Rybak IA (2025). Ionic mechanisms underlying bistability in spinal motoneurons: insights from a computational model. Front Cell Neurosci 19, 1710893.

Mousa MH & Elbasiouny SM (2021). Estimating the effects of slicing on the electrophysiological properties of spinal motoneurons under normal and disease conditions. J Neurophysiol 125, 1450–1467.

Murray KC, Nakae A, Stephens MJ, Rank M, D’Amico J, Harvey PJ, Li X, Harris RLW, Ballou EW, Anelli R, Heckman CJ, Mashimo T, Vavrek R, Sanelli L, Gorassini MA, Bennett DJ & Fouad K (2010). Recovery of motoneuron and locomotor function after spinal cord injury depends on constitutive activity in 5-HT2C receptors. Nat Med 16, 694–700.

Nascimento F, Broadhead MJ, Tetringa E, Tsape E, Zagoraiou L & Miles GB (2020). Synaptic mechanisms underlying modulation of locomotor-related motoneuron output by premotor cholinergic interneurons. Elife; DOI: 10.7554/eLife.54170.

Nascimento F, Özyurt MG, Halablab K, Bhumbra GS, Caron G, Bączyk M, Zytnicki D, Manuel M, Roselli F, Brownstone R & Beato M (2024). Spinal microcircuits go through multiphasic homeostatic compensations in a mouse model of motoneuron degeneration. Cell Rep 43, 115046.

Orssatto LBR, Borg DN, Blazevich AJ, Sakugawa RL, Shield AJ & Trajano GS (2021). Intrinsic motoneuron excitability is reduced in soleus and tibialis anterior of older adults. Geroscience 43, 2719–2735.

Özyurt MG, Ojeda-Alonso J, Beato M & Nascimento F (2022). In vitro longitudinal lumbar spinal cord preparations to study sensory and recurrent motor microcircuits of juvenile mice. J Neurophysiol 128, 711–726.

Pagiazitis JG, Delestrée N, Sowoidnich L, Sivakumar N, Simon CM, Chatzisotiriou A, Albani M & Mentis GZ (2025). Catecholaminergic dysfunction drives postural and locomotor deficits in a mouse model of spinal muscular atrophy. Cell Rep 44, 115147.

Perrier J-F, Rasmussen HB, Christensen RK & Petersen AV (2013). Modulation of the intrinsic properties of motoneurons by serotonin. Curr Pharm Des 19, 4371–4384.

Picton LD, Zhang H & Sillar KT (2017). Sodium pump regulation of locomotor control circuits. J Neurophysiol 118, 1070–1081.

Pocratsky AM, Nascimento F, Özyurt MG, White IJ, Sullivan R, O’Callaghan BJ, Smith CC, Surana S, Beato M & Brownstone RM (2023). Pathophysiology of Dyt1- dystonia in mice is mediated by spinal neural circuit dysfunction. Sci Transl Med 15, eadg3904.

Powers RK & Binder MD (2003). Persistent sodium and calcium currents in rat hypoglossal motoneurons. J Neurophysiol 89, 615–624.

Powers RK & Heckman CJ (2015). Contribution of intrinsic motoneuron properties to discharge hysteresis and its estimation based on paired motor unit recordings: a simulation study. J Neurophysiol 114, 184–198.

Powers RK, Nardelli P & Cope TC (2008). Estimation of the contribution of intrinsic currents to motoneuron firing based on paired motoneuron discharge records in the decerebrate cat. J Neurophysiol 100, 292–303.

Prinz AA, Bucher D & Marder E (2004). Similar network activity from disparate circuit parameters. Nat Neurosci 7, 1345–1352.

Quinlan KA, Schuster JE, Fu R, Siddique T & Heckman CJ (2011). Altered postnatal maturation of electrical properties in spinal motoneurons in a mouse model of amyotrophic lateral sclerosis. J Physiol 589, 2245–2260.

Reedich EJ, Genry LT, Steele PR, Mena Avila E, Dowaliby L, Drobyshevsky A, Manuel M & Quinlan KA (2023). Spinal motoneurons respond aberrantly to serotonin in a rabbit model of cerebral palsy. J Physiol 601, 4271–4289.

Revill AL, Chu NY, Ma L, LeBlancq MJ, Dickson CT & Funk GD (2019). Postnatal development of persistent inward currents in rat XII motoneurons and their modulation by serotonin, muscarine and noradrenaline. J Physiol 597, 3183–3201.

Revill AL & Fuglevand AJ (2011). Effects of persistent inward currents, accommodation, and adaptation on motor unit behavior: a simulation study. J Neurophysiol 106, 1467–1479.

Sawczuk A, Powers RK & Binder MD (1995). Spike frequency adaptation studied in hypoglossal motoneurons of the rat. J Neurophysiol 73, 1799–1810.

Schwindt PC & Crill WE (1980). Properties of a persistent inward current in normal and TEA-injected motoneurons. J Neurophysiol 43, 1700–1724.

Schwindt P & Crill WE (1977). A persistent negative resistance in cat lumbar motoneurons. Brain Res 120, 173–178.

Sharples SA, Broadhead MJ, Gray JA & Miles GB (2023). M-type potassium currents differentially affect activation of motoneuron subtypes and tune recruitment gain. J Physiol 601, 5751–5775.

Sharples SA, Koblinger K, Humphreys JM & Whelan PJ (2014). Dopamine: a parallel pathway for the modulation of spinal locomotor networks. Front Neural Circuits 8, 55.

Sharples SA & Miles GB (2021). Maturation of persistent and hyperpolarization-activated inward currents shapes the differential activation of motoneuron subtypes during postnatal development. Elife; DOI: 10.7554/eLife.71385.

Sharples SA, Nisbet SJ, Broadhead MJ, Bo Jensen D, Sorrell FL, Meehan CF & Miles GB (2025). Intrinsic mechanisms contributing to the biophysical signature of mouse gamma motoneurons. J Physiol 603, 7145–7170.

Singh S, Shevtsova NA, Yao L, Rybak IA & Dougherty KJ (2025). Properties of rhythmogenic currents in spinal Shox2 interneurons across postnatal development. J Physiol 603, 3201–3221.

Škarabot J, Beauchamp JA & Pearcey GEP (2025). Human motor unit discharge patterns reveal differences in neuromodulatory and inhibitory drive to motoneurons across contraction levels. J Neurophysiol 134, 1429–1444.

Smith CC & Brownstone RM (2020). Spinal motoneuron firing properties mature from rostral to caudal during postnatal development of the mouse. J Physiol 598, 5467–5485.

Steele PR, Cavarsan CF, Dowaliby L, Westefeld M, Katenka N, Drobyshevsky A, Gorassini MA & Quinlan KA (2020). Altered Motoneuron Properties Contribute to Motor Deficits in a Rabbit Hypoxia-Ischemia Model of Cerebral Palsy. Front Cell Neurosci 14, 69.

Stifani N (2011). Generation of Motor Neuron Diversity in the Cervical Spinal Cord.

Tiwari MN, Mohan S, Biala Y & Yaari Y (2018). Differential contributions of Ca -activated K channels and Na /K -ATPases to the generation of the slow afterhyperpolarization in CA1 pyramidal cells. Hippocampus 28, 338–357.

Udina E, D’Amico J, Bergquist AJ & Gorassini MA (2010). Amphetamine increases persistent inward currents in human motoneurons estimated from paired motor-unit activity. J Neurophysiol 103, 1295–1303.

Vandenberk MS & Kalmar JM (2014). An evaluation of paired motor unit estimates of persistent inward current in human motoneurons. J Neurophysiol 111, 1877–1884.

Verneuil J, Brocard C, Trouplin V, Villard L, Peyronnet-Roux J & Brocard F (2020). The M-current works in tandem with the persistent sodium current to set the speed of locomotion. PLoS Biol 18, e3000738.

Vinay L, Brocard F, Pflieger JF, Simeoni-Alias J & Clarac F (2000). Perinatal development of lumbar motoneurons and their inputs in the rat. Brain Res Bull 53, 635–647.

